# Improvement in Orchard Nutrient Status and Productivity Through Integrated Nutrient Management in Mango (*Mangifera indica* L.) Under Medium-Density Planting

**DOI:** 10.64898/2025.12.30.697138

**Authors:** Kuldeep, Ashok Kumar Singh, Amit Bhatnagar, Omveer Singh, Satya Pratap Pachauri, Shailesh Chandra Shankhdhar, Hem Ch. Joshi, Manoj Kumar Bhatt

## Abstract

Balanced nutrient management is essential for sustaining productivity and soil health in perennial fruit crops. A two-year field study (2022-2024) was conducted to evaluate the effects of integrating soil and foliar micronutrient applications with reduced recommended doses of fertilizers (RDF) on vegetative growth, nutrient status, yield, fruit quality, and economic performance of mango (*Mangifera indica* L.) cv. Dashehari under medium-density planting. Nine nutrient modules combining 25-100% RDF with soil- and foliar-applied Fe, Zn, B, Cu, and Ca were compared using ANOVA, PCA, and PLS-SEM. Integrated treatments significantly enhanced canopy height, spread, and volume, with the 75% RDF + two foliar micronutrient sprays producing the greatest vegetative growth. Yield attributes, fruit size, retention, and shelf life improved markedly under micronutrient-enriched treatments. The highest fruit yield and benefit-cost ratio was obtained with 75% RDF + two foliar sprays, outperforming the conventional 100% RDF. Soil pH and EC were moderated, and organic carbon improved slightly under integrated nutrient management. Soil and leaf macro- and micronutrient concentrations were consistently higher with combined soil-foliar approaches. PCA clustered high-performing nutrient modules with yield and canopy variables, while PLS-SEM identified plant nutrient status as the strongest predictor of fruit yield. Overall, integrating micronutrient supplementation with 75% RDF improved nutrient-use efficiency, productivity, and economic returns while reducing dependence on full-dose fertilizers. The study highlights 75% RDF + foliar micronutrient sprays as an effective and sustainable nutrient management strategy for enhancing yield and orchard performance in mango.

**Graphical abstract:** 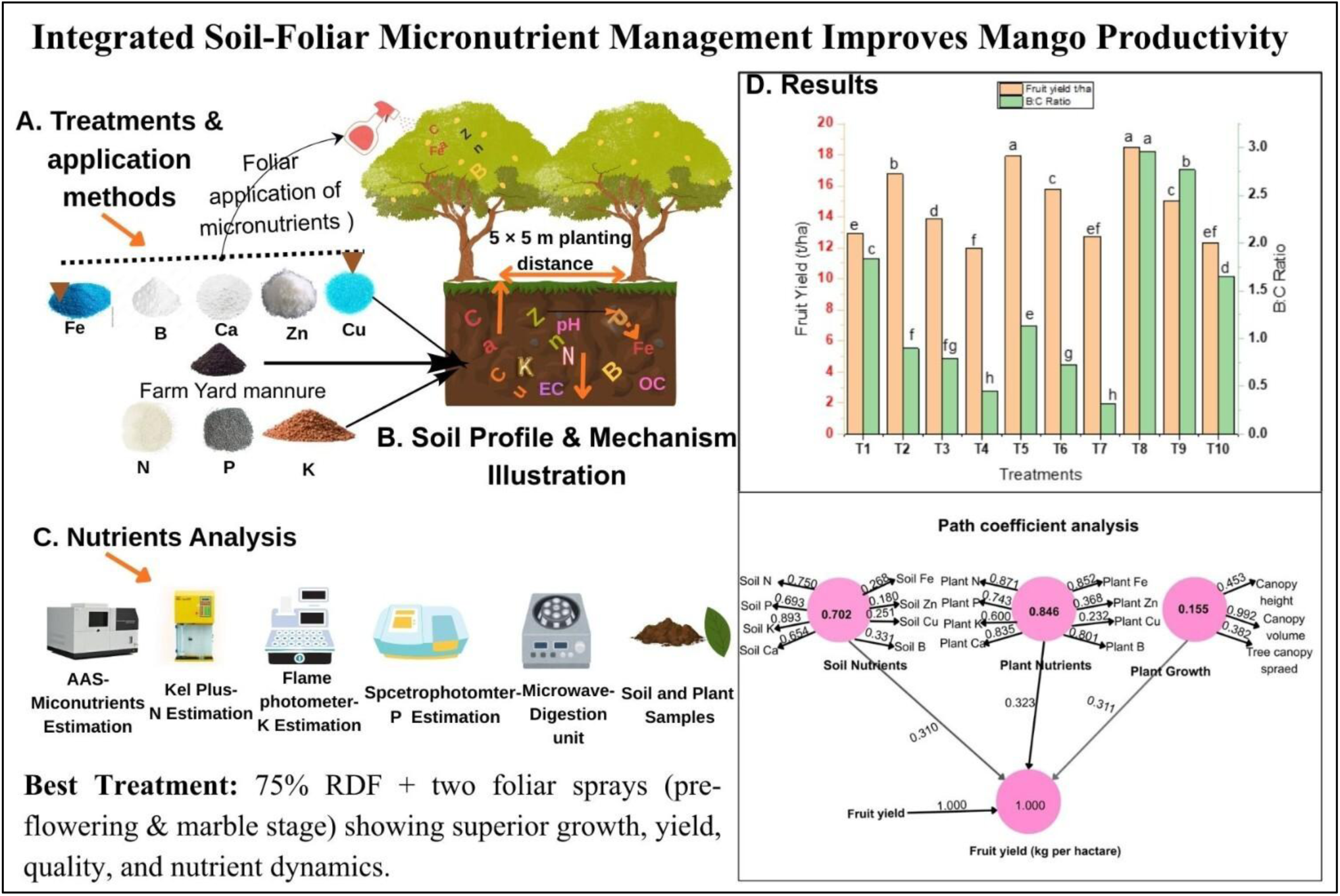

## 1. Introduction

Mango (*Mangifera indica* L.) is a major perennial fruit crop of the tropics and subtropics and the most widely cultivated species of the family Anacardiaceae. Its domestication in the Indian subcontinent dates back more than four millennia, reflecting its long-standing association with human societies (Kaur 2017; Pandey et al. 2025). Believed to have originated in the Indo-Burma region, mango exhibits broad adaptability to diverse agroclimatic conditions and thrives across a range of soil types, temperatures, and rainfall regimes (Singh et al. 2016; Nasron et al. 2021). Cytologically, M. indica is diploid (2n = 40), and the genus Mangifera comprises over 26 related species distributed mainly in Southeast Asia, several of which bear edible fruits (Yang et al. 2022; Hussain et al. 2021). Recognized globally as the “King of Fruits,” mango is valued for its distinctive aroma, rich biochemical composition, and balanced flavor (Maldonado-Celis et al. 2019; Peng et al. 2022). Mango fruits are nutritionally essential, providing vitamin A (≈4800 IU 100 g⁻¹), vitamin C, B-complex vitamins, malic acid, proteins, sugars, fats, and important minerals such as calcium and iron (Lebaka et al. 2021; Hassan et al. 2022, Lingwan & Yadav 2025). India cultivates mango on nearly 2.4 million hectares, producing about 22.7 million tonnes annually, with an average productivity of 9.5 t ha⁻¹ (NHB 2024-2025). However, productivity remains below potential due to suboptimal orchard management and soil fertility constraints. Traditional orchards established with seedling rootstocks and wide spacing have low tree density and limited productivity. Increasing land pressure, urbanization, and resource constraints have accelerated the adoption of medium- and high-density planting systems, which improve land-use efficiency and yield potential (Hahn et al. 2022; Lingwan et al., 2023). Concurrent advances in canopy management, fertigation, and irrigation scheduling have further supported the expansion of these intensive systems (Tripathi et al. 2020; Singh et al. 2001). These approaches enhance productivity per unit land and water and provide viable options for sustaining fruit production under changing climatic and resource scenarios (Dalvi et al. 2010; Singh et al. 2012, Lingwan et al., 2023). Under intensified planting, efficient nutrient management becomes critical for maintaining orchard productivity and long-term soil health. Perennial fruit trees continuously remove substantial quantities of nutrients through vegetative growth and fruit harvest, necessitating regular replenishment to prevent soil fertility decline (Lather et al. 2021; Rezende et al. 2024, Koley et al.,2025). Although inorganic fertilizers are widely used due to their rapid availability, imbalanced or excessive application, particularly of NPK, can reduce nutrient-use efficiency and induce soil nutrient imbalances (Xin et al. 2025; Chen et al. 2022). Long-term fertilizer use can alter soil chemical, physical, and biological properties; nitrogen may cause acidification, while phosphorus can undergo fixation, limiting the availability of associated nutrients (Wang et al. 2022; Khan et al. 2025, Yadav et al.,2019a). Over time, disparities between nutrient addition and removal can result in depletion of secondary and micronutrients (Jones et al. 2013, Yadav et al., 2019b), thereby reducing soil health and increasing environmental risks (Vitousek et al. 2009, Jain et al.,2025). Most long-term nutrient studies have focused on annual crops, while perennial orchard systems characterized by deeper root systems and internal nutrient recycling remain comparatively understudied (Jayasinghege et al. 2024, Lingwan et al.,2025). In mango, although several investigations have reported the effects of NPK and micronutrient supplementation, these studies are generally short-term, emphasize yield alone, and often overlook the interactions among soil fertility, nutrient dynamics, and economic performance (Ravikiran 2018; Srivastava and Hu 2019; Ahmad et al. 2018; Chaudhari and Singh 2019, Shagun et al., 2024). Research on nutrient behavior in medium-density orchards is particularly limited, and advanced multivariate approaches capable of unraveling complex soil-plant-yield relationships remain largely unexplored. To address these gaps, the present study evaluates the long-term effects of integrating reduced recommended fertilizer doses (RDF) with soil and foliar micronutrient applications in a medium-density mango orchard. The study examines impacts on soil fertility, plant nutrient status, yield, and economic returns. Principal Component Analysis (PCA) and Partial Least Squares Structural Equation Modeling (PLS-SEM) were employed to analyze the causal relationships between nutrient pools and productivity. We hypothesize that integrating reduced RDF with targeted micronutrient supplementation will enhance nutrient-use efficiency, improve soil fertility, and increase yield and profitability, and that multivariate modeling will reveal positive causal pathways linking optimized nutrient pools with improved productivity in mango orchards.

## 2. Materials and Methods

### 2.1 Orchard site and experimental period

The study was conducted during 2023-2024 at the Horticulture Research Centre, Patharchatta, GBPUA&T, Pantnagar, Uttarakhand (≈29.5°N, 79.3°E; 244 m a.s.l.; **Fig.1**). The experiment used a medium-density mango (*Mangifera indica* L. cv. Dashehari) orchard of uniform 13-year-old trees planted at 5 × 5 m spacing (400 trees ha⁻¹; **Fig.2**).

**Figure 1.**
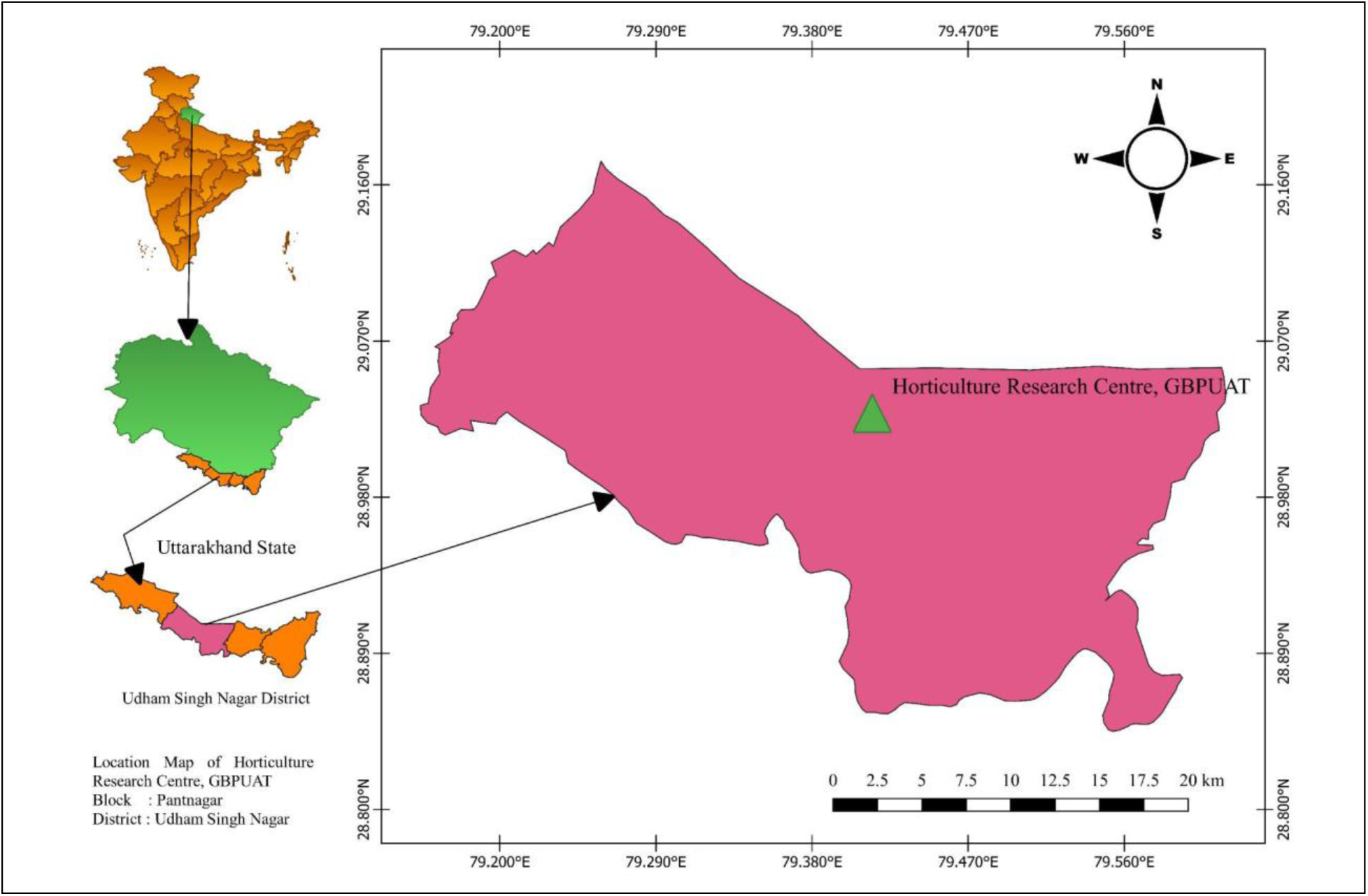
**Experimental Research Unit at HRC, Pattarchatta, U.S. Nagar, Uttarakhand (India).**

**Figure 2.**
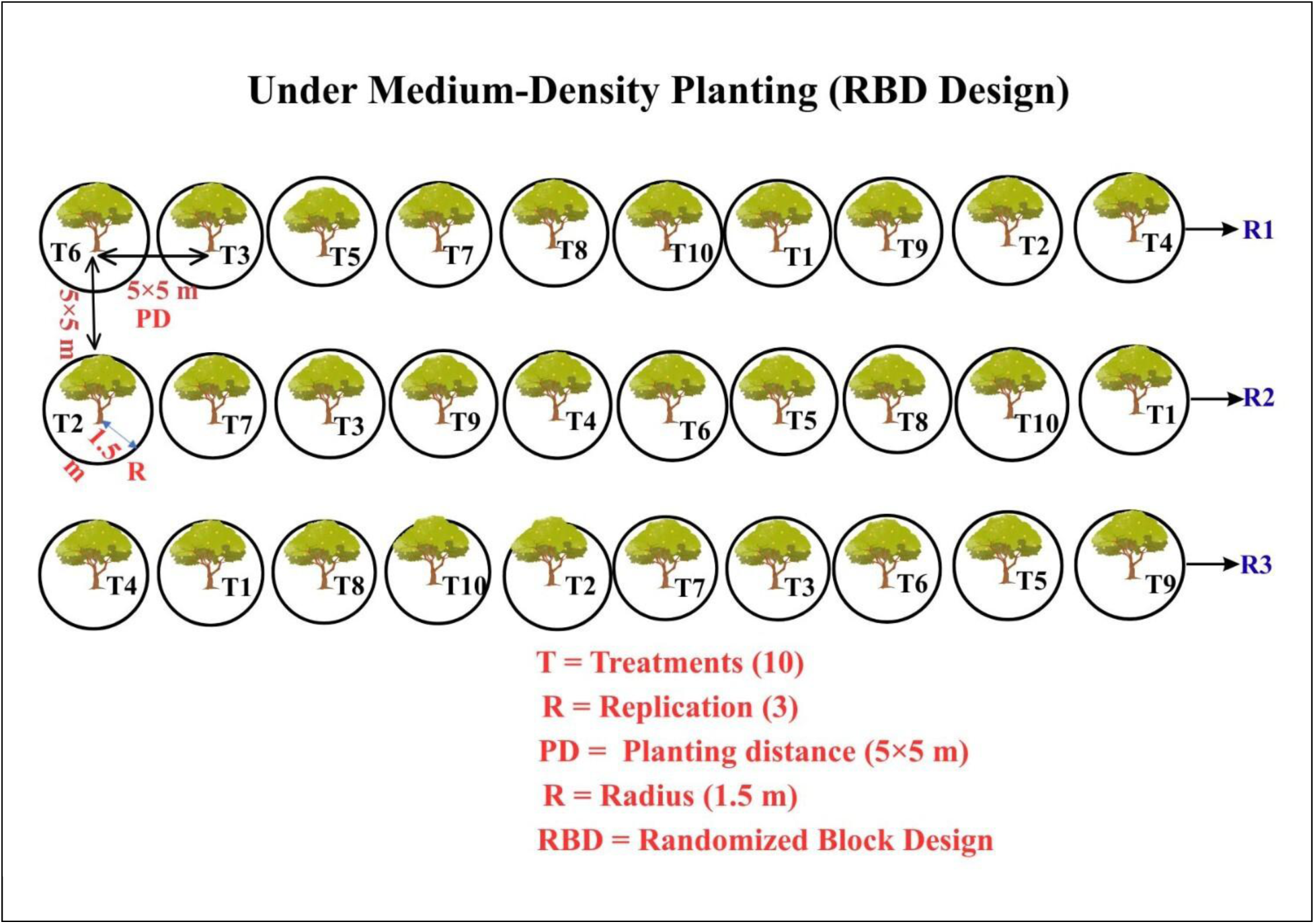
**Planting layout of mango (*Mangifera indica* L.) cv. Dashehari under INM practices in medium-density planting. T1:**100% RDF alone, **T2:** 75% RDF + soil micronutrients + one foliar spray of micronutrients, **T3:** 50% RDF + soil micronutrients + one foliar spray of micronutrients, **T4:** 25% RDF + soil micronutrients + one foliar spray of micronutrients, **T5:** 75% RDF + soil micronutrients + two foliar spray of micronutrients, **T6:** 50% RDF + soil micronutrients + two foliar spray of micronutrients, **T7:** 25% RDF + soil micronutrients + two foliar spray of micronutrients, **T8:** 75% RDF + two foliar spray of micronutrients, **T9:** 50% RDF + two foliar spray of micronutrients, **T10:** 25% RDF+ two foliar spray of micronutrients.

### 2.2 Soil and climate

The site lies in the Tarai agro-ecological zone with a sub-humid subtropical climate. The experimental soil is silty clay loam (Patharchatta Series II, Mollisol). Baseline soil physical-chemical properties (pH, EC, organic carbon, available N, P, K, and extractable micronutrients) were determined before the start of the experiment (**Table 1 for initial status**).

**Table 1.** Required/Optimum level of nutrients in the mango orchard soil and leaves.

| S. No. | Soil properties and nutrient parameters for soil & leaves | Optimum level of nutrients in the soil & leaves of the mango orchard |
| --- | --- | --- |
| **Soil nutrient status** |  |  |
| 1. | pH (1:2- Soil:Water) | 5.5-7.0 |
| 2. | Organic carbon (%) | 0.51-0.75 |
| 3. | EC (dS m<sup>-1</sup>) | <0.2 |
| 4. | Available nitrogen (kg ha<sup>-1</sup>) | 280-560 |
| 5. | Available phosphorus (kg ha<sup>-1</sup>) | 22-54 |
| 6. | Available potassium (kg ha<sup>-1</sup>) | 123-293 |
| 7. | Fe (mg kg<sup>-1</sup>) | 4.8-8.7 |
| 8. | Zn (mg kg<sup>-1</sup>) | 0.5-1.0 |
| 9. | Cu (mg kg<sup>-1</sup>) | 0.2 -0.4 |
| 10. | B (mg kg<sup>-1</sup>) | >0.5 |
| 11. | Ca (mg kg<sup>-1</sup>) | >1000 |
| **Plant nutrient status** |  |  |
| 12. | Total nitrogen (%) | 1.2-1.4 |
| 13. | Total phosphorus (%) | 0.08-0.16 |
| 14. | Total potassium (%) | 0.5-1.0 |
| 15. | Fe (ppm) | 50-200 |
| 16. | Zn (ppm) | 20-40 |
| 17. | Cu (ppm) | 10-50 |
| 18. | B (ppm) | 50-100 |
| 19. | Ca (%) | 2.0-3.5 |

**Sources:** Adak and Panday (2020), QDAF (2015), Quaggio (1996), Raja *et al.* (2005)

### 2.3 Experimental design and treatments

The experiment was laid out in a randomized block design (RBD) with 10 treatments and three replications. Treatments evaluated combinations of full, reduced (75%, 50%, 25%) recommended fertilizer dose (RDF = 1000 g N + 750 g P₂O₅ + 1000 g K₂O + 50 kg FYM tree⁻¹ year⁻¹) and soil-applied micronutrients (Fe, Cu, Zn each 100 g tree⁻¹ and B 50 g tree⁻¹), together with one or two foliar sprays of micronutrients. In brief treatment details:

- T1 (Control): 100% RDF applied in the basin after harvest.
- T2-T4: 75%, 50%, 25% RDF, respectively, + soil micronutrients (Fe, Cu, Zn @100 g each + B @50 g tree⁻¹) + one foliar spray (Fe, Ca, Zn each 0.50% + B- 0.10%) (just before flowering and marble stages as a single event).
- T5-T7: 75%, 50%, 25% RDF, respectively, + soil micronutrients (as above) + two foliar sprays (same foliar composition; applied just before flowering and at marble stage).
- T8-T10: 75%, 50%, 25% RDF, respectively, without soil micronutrients but with two foliar sprays (just before flowering and at marble stage).

(Note: Full treatment details are summarized in Table X: Supplementary materials)

### 2.4 Orchard management and fertilizer application

Basins (radius: 1.5 m from trunk) were prepared around each tree. Farmyard manure was applied in October. P, K, and micronutrient basal doses were incorporated into the basins in November. Nitrogen was applied in two equal splits: first immediately after flowering and the second at the mustard-to-pea stage of fruit development. Foliar sprays (10 L water tree⁻¹ per spray) were prepared by dissolving micronutrient salts and applied with a tractor sprayer at the timings (just before flowering and at marble stage). Standard cultural practices (irrigation, weeding, pruning, pest and disease control) were followed uniformly across treatments. Conceptual experimental workflow for the integrated nutrient management study in medium-density mango orchards is presented in **(Fig.3),** and the methods and timings of nutrient application are illustrated in (**Fig. 4).**

**Figure 3.**
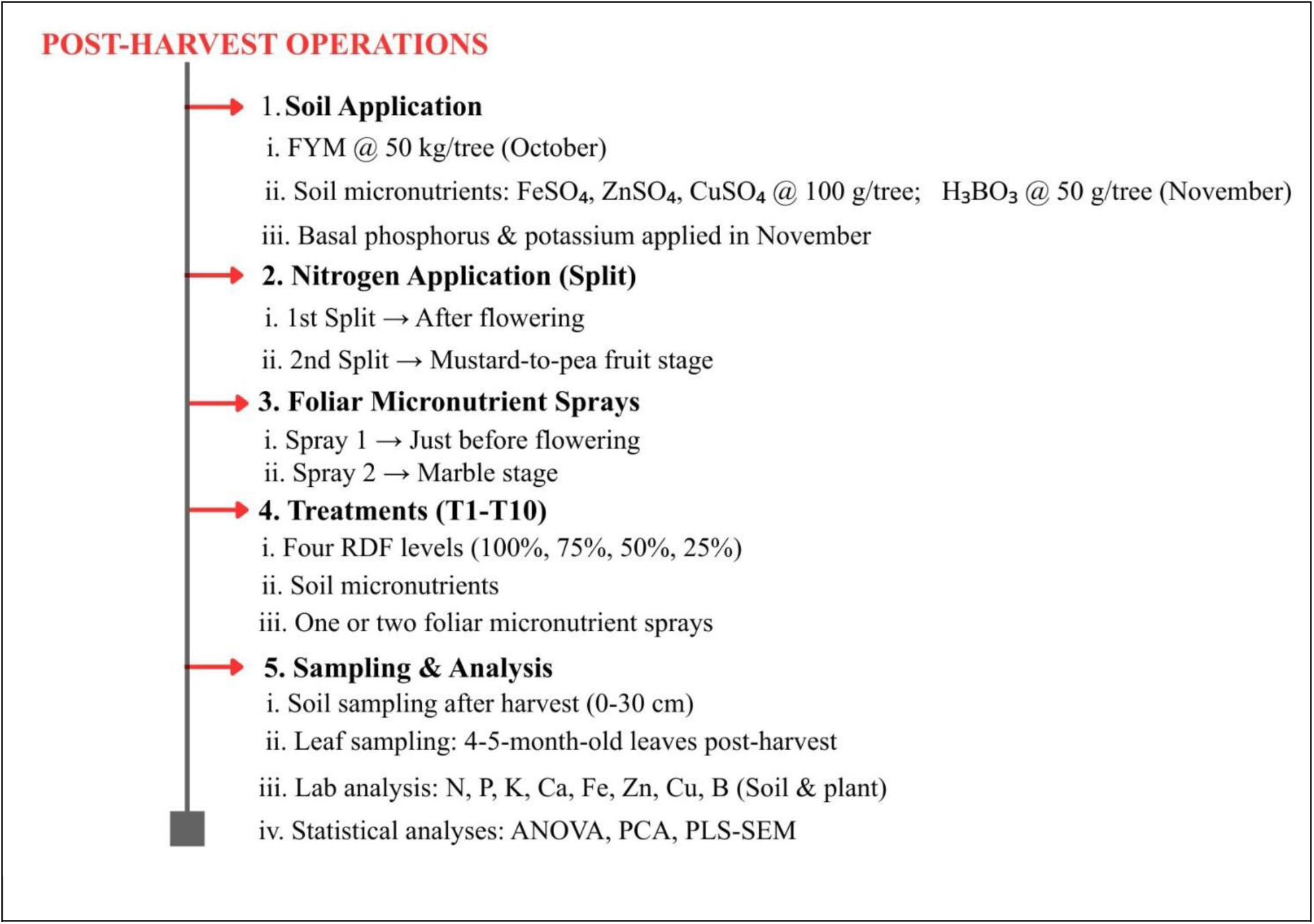
**Conceptual experimental workflow for the integrated nutrient management study in medium-density mango orchards.**

**Figure 4.**
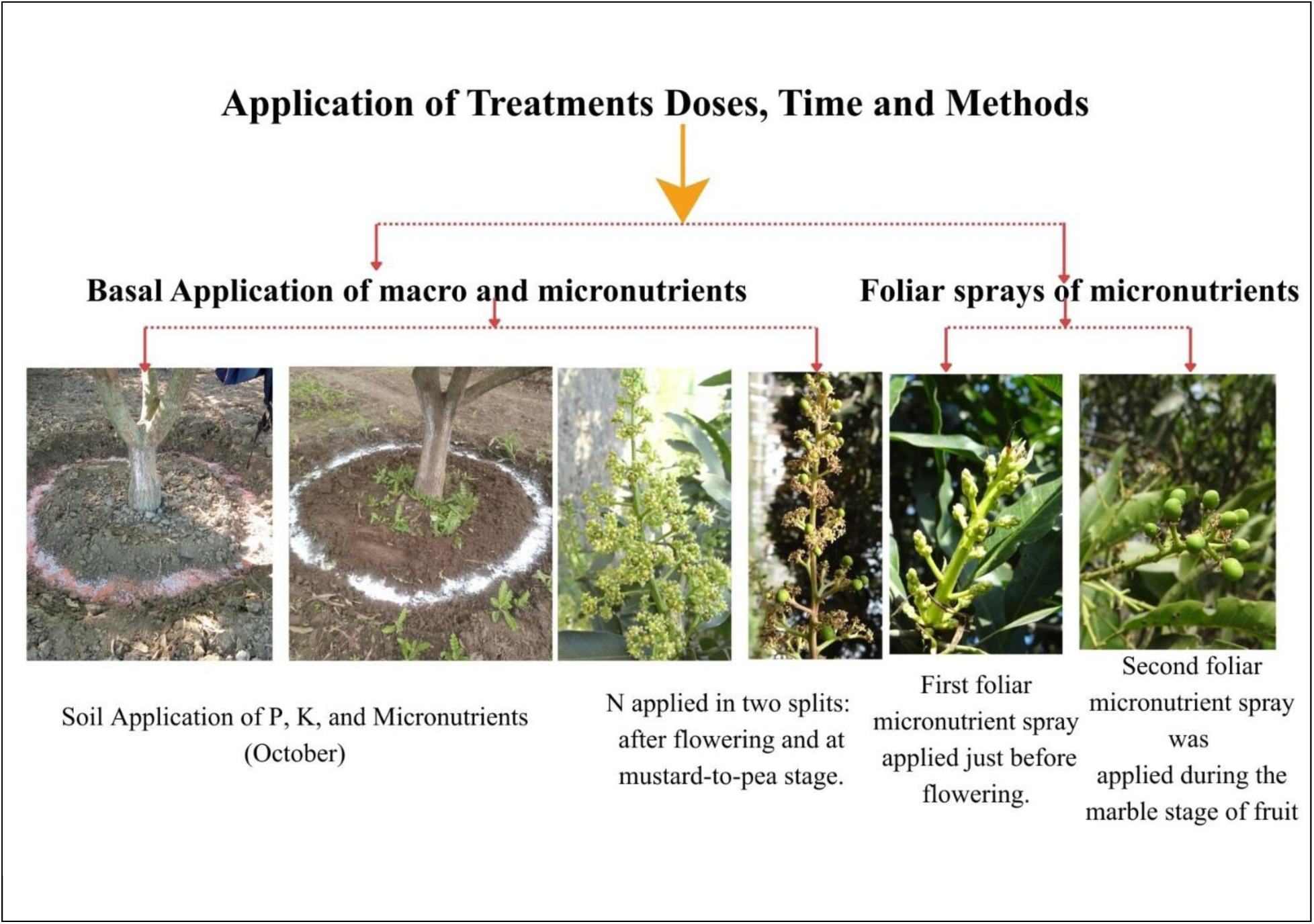
Overview of the doses of fertilizer, time, and application methods (*Mangifera indica* L.) cv. Dashehari for mango Crops under medium density planting. T1:100% RDF alone, **T2:** 75% RDF + soil micronutrients + one foliar spray of micronutrients, **T3:** 50% RDF + soil micronutrients + one foliar spray of micronutrients, **T4:** 25% RDF + soil micronutrients + one foliar spray of micronutrients, **T5:** 75% RDF + soil micronutrients + two foliar spray of micronutrients, **T6:** 50% RDF + soil micronutrients + two foliar spray of micronutrients, **T7:** 25% RDF + soil micronutrients + two foliar spray of micronutrients, **T8:** 75% RDF + two foliar spray of micronutrients, **T9:** 50% RDF + two foliar spray of micronutrients, **T10:** 25% RDF+ two foliar spray of micronutrients.

### 2.5 Sampling and laboratory analysis

#### 2.5.1 Soil sampling

Soil samples were collected from each replication after harvest in both years from the 0-30 cm depth using a post-hole auger. Samples were air-dried, sieved (<2 mm), and stored for analysis. Soil pH and electrical conductivity (EC) were determined in 1:2.5 soil: water extracts. Organic carbon was estimated by Walkley-Black wet oxidation. Available N was measured by the alkaline permanganate method; available P by Olsen’s bicarbonate extraction and colorimetry; available K by ammonium acetate extraction and flame photometry. DTPA extraction (pH 7.3) was used for available Fe, Zn, and Cu, and concentrations were measured by atomic absorption spectrophotometry (AAS) (Pant et al., 2023). Boron was estimated by the hot CaCl₂ extractable method and quantified spectrophotometrically. An overview of soil sample collection (0-30 cm depth) is presented in **Fig. 5**.

**Figure 5.**
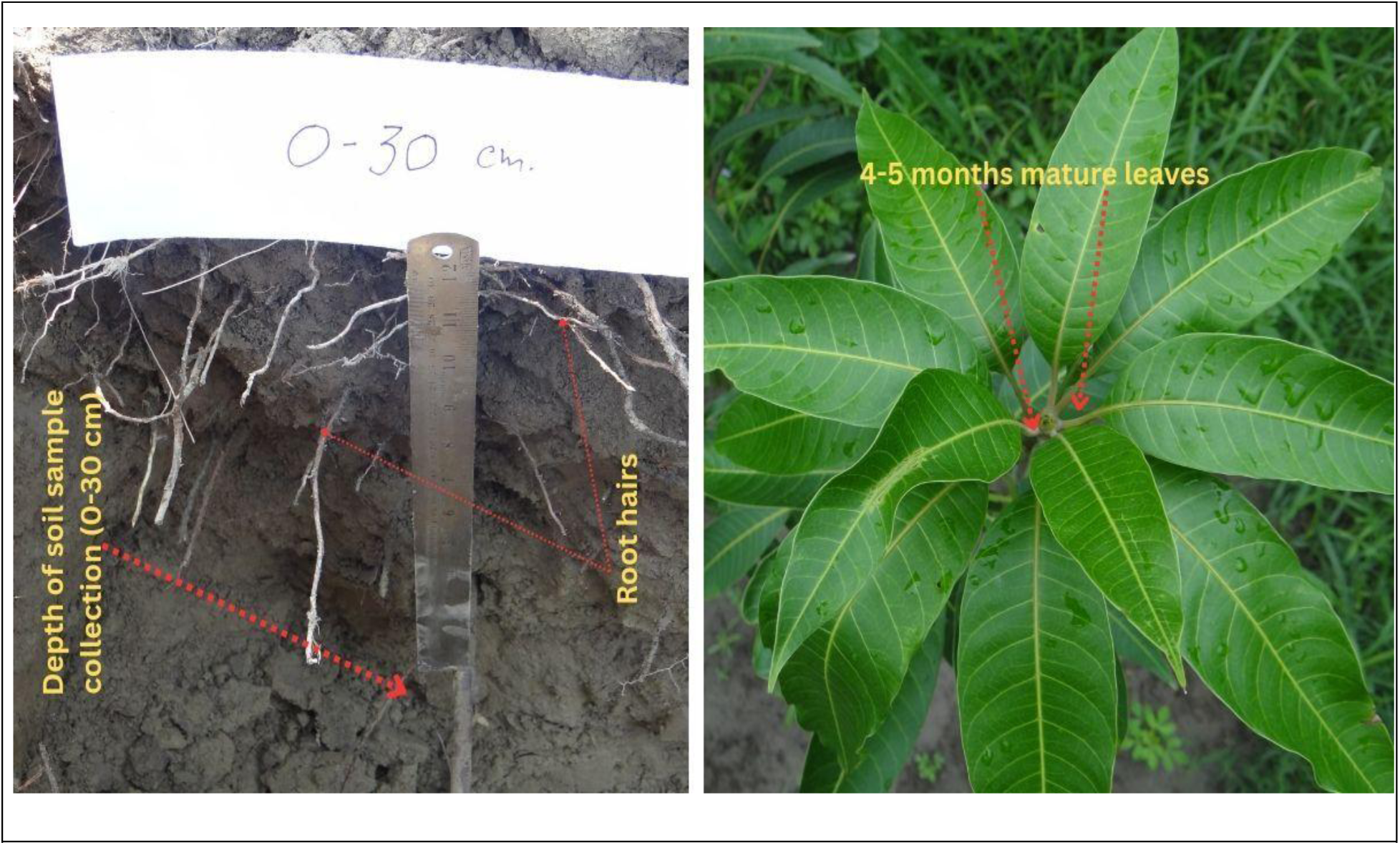
**Overview of soil sample collection (0-30 cm depth) and plant sample collection from 4-5-month-old mature mango leaves.**

#### 2.5.2 Plant sampling

Mature (4-5-month-old) fully expanded leaves were collected after harvest from each tree for nutrient analysis **Fig. 5**. Leaves were washed, oven-dried at 65 ± 1 °C to constant weight, ground, and sieved (<2 mm). Plant tissue digestion was performed using di-acid (HNO₃:HClO₄) digestion. Total N was determined by the Kjeldahl method, P by colorimetry (vanadate–molybdate), K by flame photometry, and Fe, Zn, Cu, and Ca by AAS. Boron in plant tissue was determined after dry-ashing by the Azomethine-H method.

### 2.6 Growth, yield, quality, and benefit: cost ratio measurements

Tree height (from first branch to apex), canopy spread (N-S and E-W), canopy volume (Westwood & Roberts, 1970), and their percentage increases were recorded at the start and end of the experimental period. Fruit set, fruit drop, and fruit retention were recorded on tagged panicles. At harvest, fruit number per tree and mean fruit weight (mean of 20-25 fruits per tree) were used to calculate yield per tree (kg tree⁻¹) and extrapolated to t ha⁻¹. Yield efficiency was calculated as yield per unit canopy volume (kg m⁻³). Fruit dimensions (length, width) and fruit weight were measured on representative samples using digital calipers and a balance. Shelf life was assessed at ambient temperature (25 ± 2 °C) by recording days until acceptable edible ripeness. The B: C ratio was determined to assess the economic efficiency of each treatment, showing the returns per rupee invested.

### 2.7 Multivariate analyses and statistics

Data for two consecutive years were pooled after testing for homogeneity of error variance. Treatment effects were tested by analysis of variance (ANOVA) appropriate for an RBD; when significant, means were separated using Tukey’s HSD at P ≤ 0.05. The Persian correlation coefficient was calculated to quantify the strength and direction of associations among soil, plant, and yield variables, providing a robust non-parametric measure suitable for non-normal or heterogeneous datasets. Principal Component Analysis (PCA) was used to identify major patterns among soil, plant, and yield variables (all variables standardized before analysis) (Shree et al., 2019; Lingwan et al., 2022). Partial Least Squares Structural Equation Modeling (PLS-SEM) was performed to evaluate causal pathways among latent constructs representing Plant Growth, Plant Nutrients, Soil Nutrients, and Fruit Yield. PCA was run in OriginPro; PLS-SEM and path modelling were implemented in SmartPLS (version 4.1.1.5). Model quality was assessed using standard metrics (R², composite reliability, AVE, discriminant validity, VIF, and SRMR).

## 3. Results

### 3.1. Vegetative growth

Canopy growth parameters of mango cv. Dashehari responded significantly to nutrient management treatments during 2022-2023 and 2023-2024 **(Table 2).** Canopy height, spread and volume varied markedly across treatments. The treatment receiving 75% RDF with two foliar micronutrient sprays recorded the maximum canopy height, whereas the 100% RDF treatment consistently produced the lowest values. Canopy spread was greatest under the integrated module of 50% RDF + soil + foliar micronutrients, while the minimum spread occurred in the treatment receiving 50% RDF with micronutrients but fewer foliar sprays. Canopy volume differed significantly, with 75% RDF + two foliar sprays producing the largest volume, statistically superior to all other treatments. Treatments integrating reduced RDF (75% and 50%) with micronutrients produced significantly larger canopy volumes than the sole 100% RDF treatment, which recorded the lowest canopy volume.

**Table 2.**
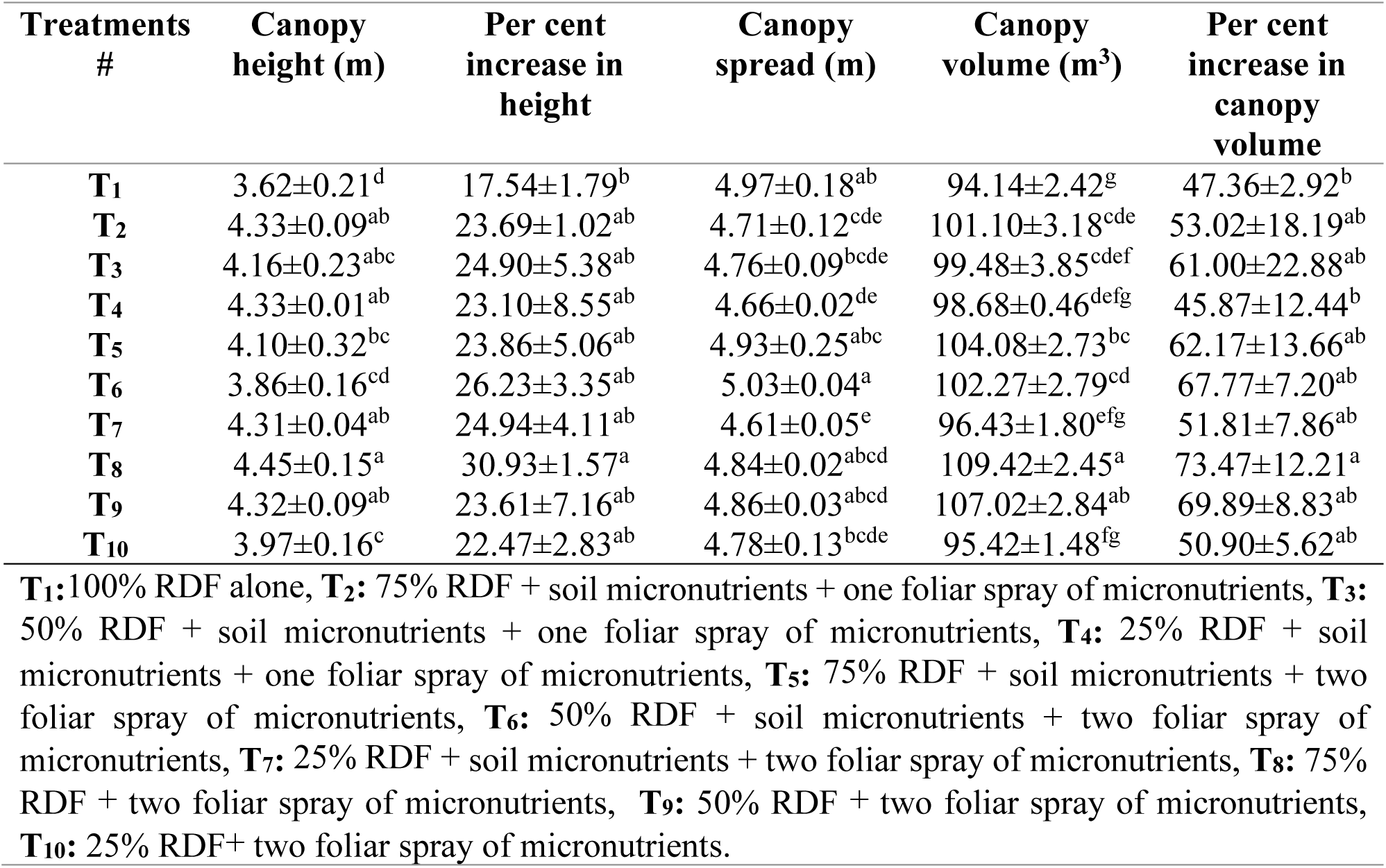
Influence of macro and micronutrients on the vegetative growth of mango cv. Dashehari under medium density planting (**Mean of two years**).

### 3.2. Yield attributes, shelf life, and economic returns

Yield attributes, fruit quality and economic traits were significantly influenced by soil and foliar micronutrient treatments (Table 3; Fig. 6). The number of fruits per panicle was highest under 50% RDF + two foliar sprays, comparable with 75% RDF + two sprays and 25% RDF + sprays, while the control (100% RDF) produced the fewest fruits. Fruit retention at maturity was highest in 75% RDF + soil + foliar micronutrients, whereas reduced RDF treatments with limited micronutrient inputs showed the lowest retention. Fruit drop followed an inverse trend.

**Figure 6.**
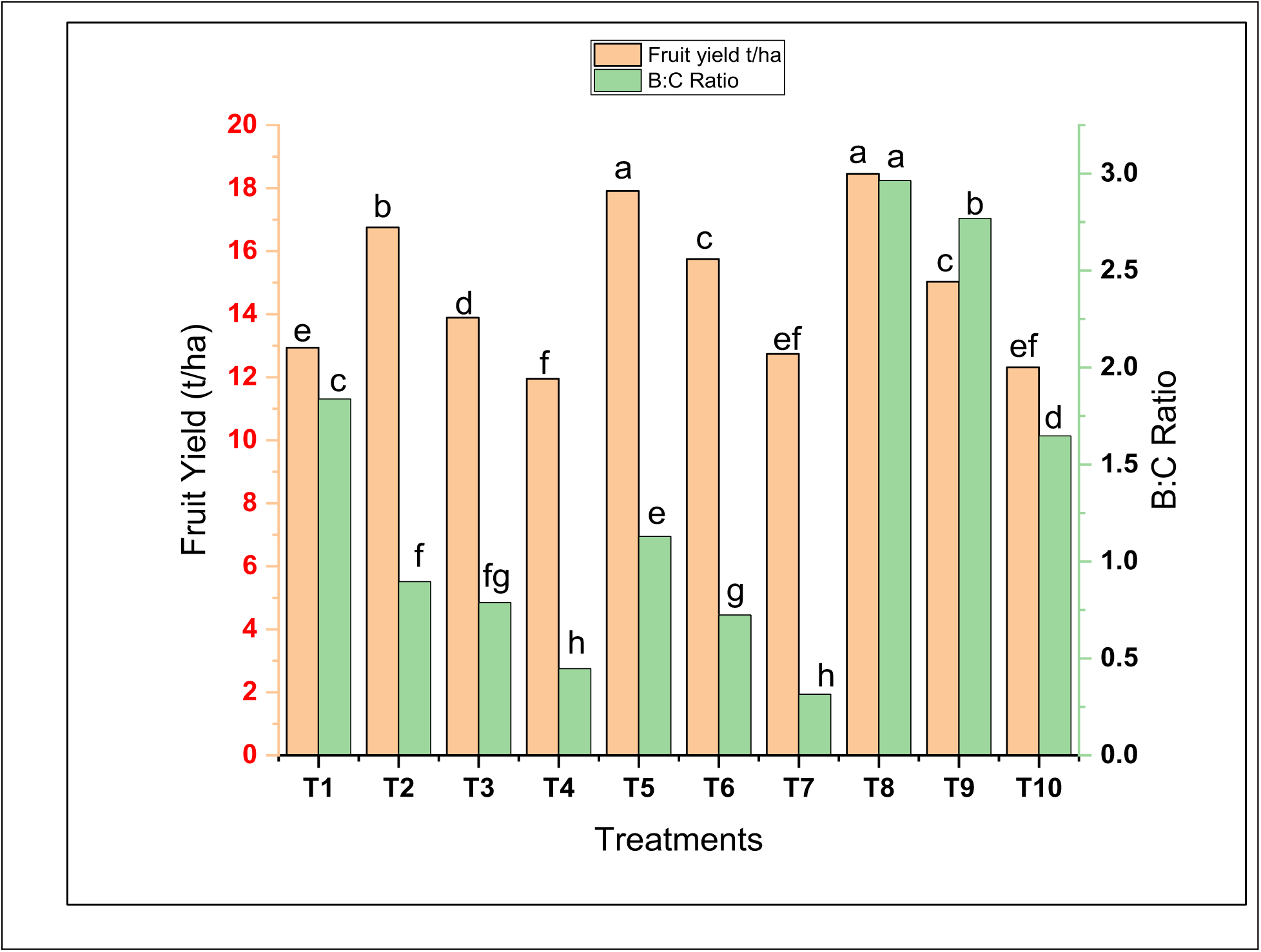
Influence of INM practices on yield and B:C Ratio of mango (*Mangifera indica* L.) cv. Dashehari under medium-density planting (Mean of two years). T1:100% RDF alone, **T2:** 75% RDF + soil micronutrients + one foliar spray of micronutrients, **T3:** 50% RDF + soil micronutrients + one foliar spray of micronutrients, **T4:** 25% RDF + soil micronutrients + one foliar spray of micronutrients, **T5:** 75% RDF + soil micronutrients + two foliar spray of micronutrients, **T6:** 50% RDF + soil micronutrients + two foliar spray of micronutrients, **T7:** 25% RDF + soil micronutrients + two foliar spray of micronutrients, **T8:** 75% RDF + two foliar spray of micronutrients, **T9:** 50% RDF + two foliar spray of micronutrients, **T10:** 25% RDF+ two foliar spray of micronutrients.

**Table 3.**
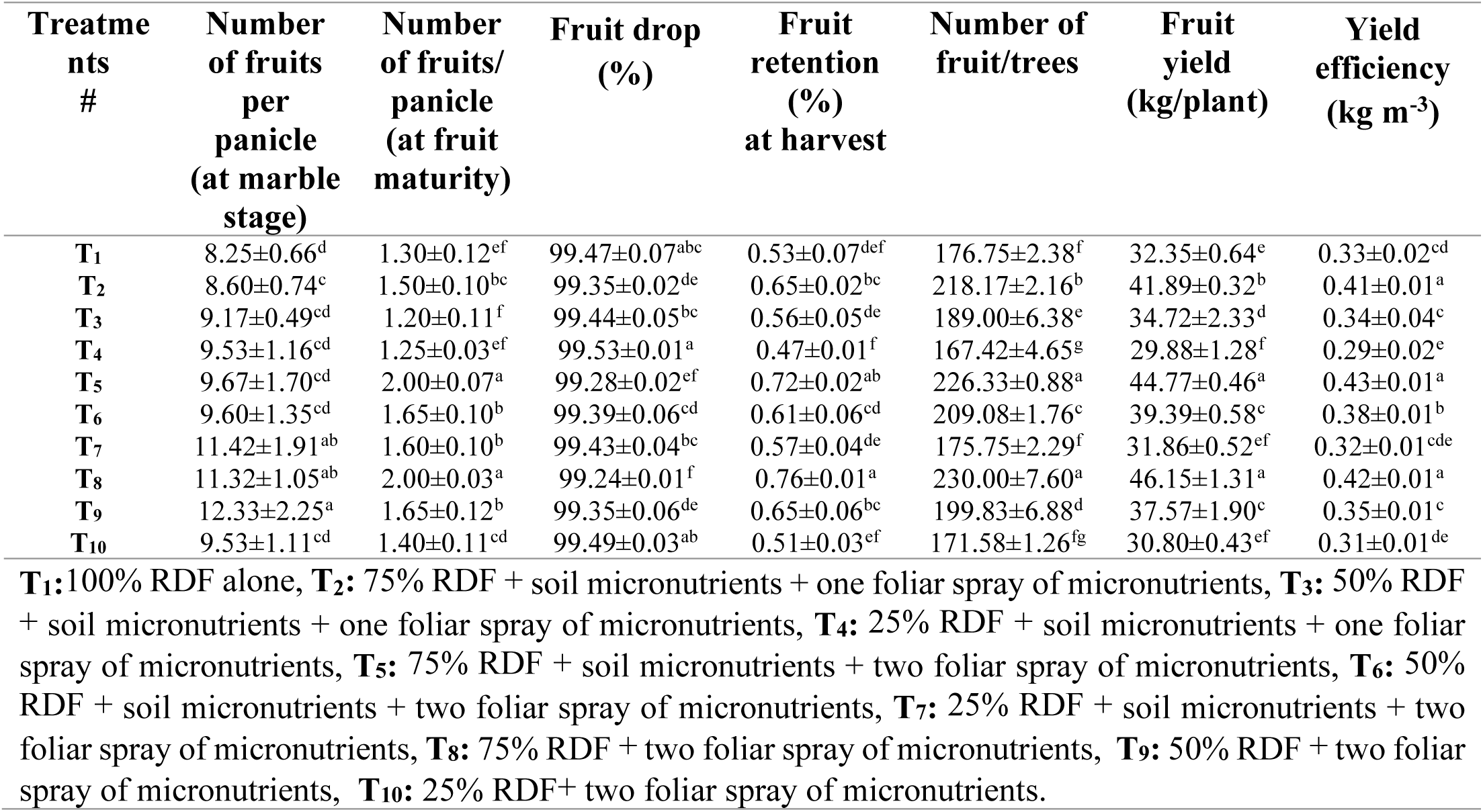
Influence of macro and micronutrients on the yield attributes of mango cv. Dashehari under medium density planting (**Mean of two years**).

Fruit yield per tree and per hectare was significantly affected. The treatment 75% RDF + two foliar sprays produced the highest yield, followed closely by 75% RDF + soil + foliar application. The lowest yield occurred in 25% RDF. Yield efficiency also increased under integrated nutrient treatments. Fruit size attributes (length, width, weight) were highest where foliar micronutrient sprays were applied. Shelf life increased notably in foliar-spray treatments. Economic analysis indicated the highest B:C ratio under 75% RDF + foliar spray, followed by 50% RDF + foliar spray.

### 3.3. Soil pH, organic carbon and electrical conductivity

Soil chemical properties varied significantly among treatments **(Table 4).** Soil pH was highest under 100% RDF and lowest under integrated treatments combining reduced RDF with micronutrient applications. Organic carbon showed slight variations, with the highest values observed in 75% RDF + soil + foliar micronutrients, and the lowest under reduced RDF with limited micronutrients. Soil electrical conductivity varied within a narrow range but was significantly higher in the 100% RDF treatment. The lowest EC was recorded in the 25% RDF + soil + foliar micronutrients treatment.

**Table 4.**
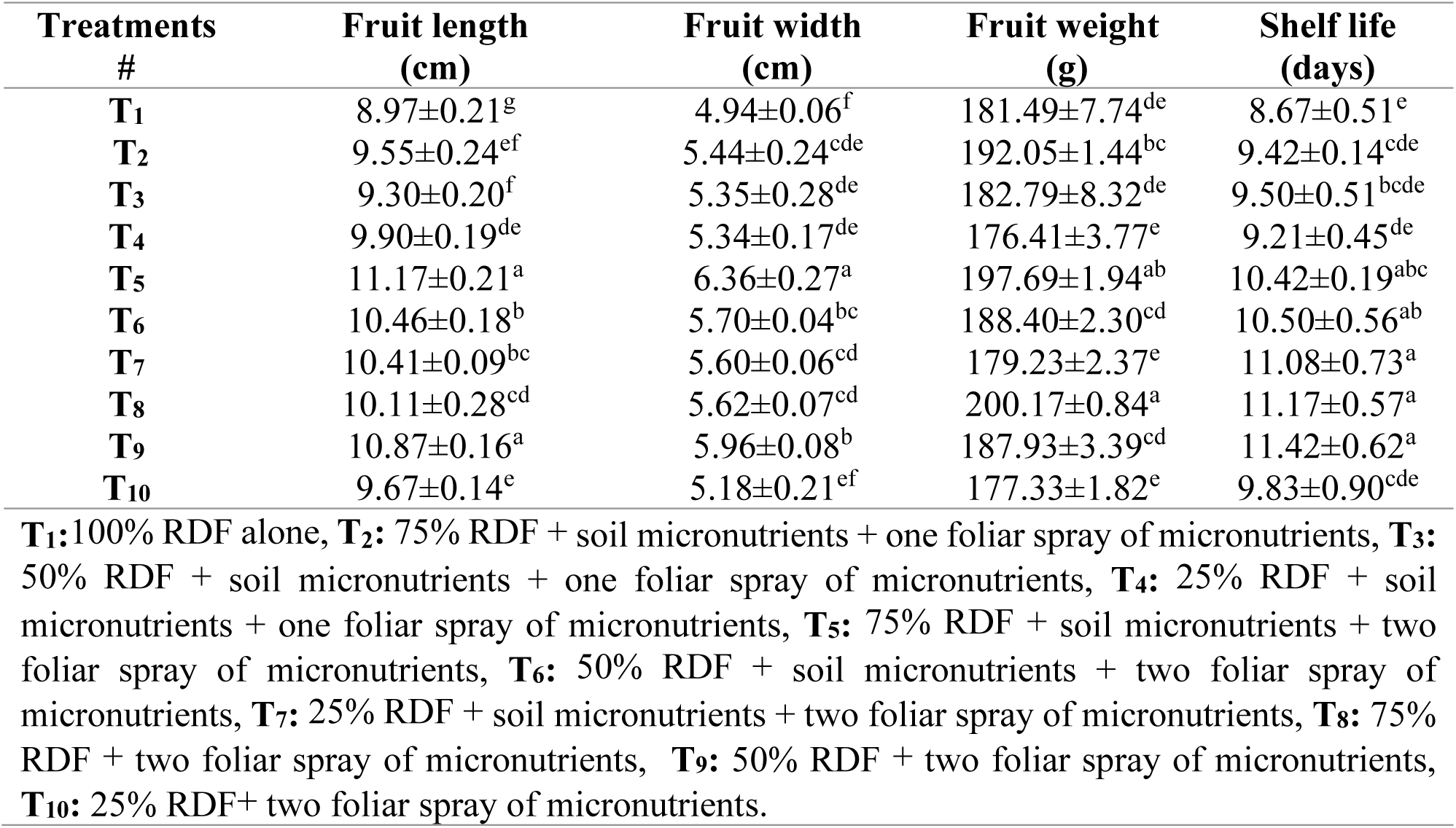
Influence of macro and micronutrients on fruit size, weight, and shelf life of mango cv. Dashehari under medium density planting (**Mean of two years**).

| **Treatments**<br>**#** | **Fruit length**<br>**(cm)** | **Fruit width**<br>**(cm)** | **Fruit weight**<br>**(g)** | **Shelf life**<br>**(days)** |
| --- | --- | --- | --- | --- |
| **T<sub>1</sub>** | 8.97±0.21<sup>g</sup> | 4.94±0.06<sup>f</sup> | 181.49±7.74<sup>de</sup> | 8.67±0.51<sup>e</sup> |
| **T<sub>2</sub>** | 9.55±0.24<sup>ef</sup> | 5.44±0.24<sup>cde</sup> | 192.05±1.44<sup>bc</sup> | 9.42±0.14<sup>cde</sup> |
| **T<sub>3</sub>** | 9.30±0.20<sup>f</sup> | 5.35±0.28<sup>de</sup> | 182.79±8.32<sup>de</sup> | 9.50±0.51<sup>bcde</sup> |
| **T<sub>4</sub>** | 9.90±0.19<sup>de</sup> | 5.34±0.17<sup>de</sup> | 176.41±3.77<sup>e</sup> | 9.21±0.45<sup>de</sup> |
| **T<sub>5</sub>** | 11.17±0.21<sup>a</sup> | 6.36±0.27<sup>a</sup> | 197.69±1.94<sup>ab</sup> | 10.42±0.19<sup>abc</sup> |
| **T<sub>6</sub>** | 10.46±0.18<sup>b</sup> | 5.70±0.04<sup>bc</sup> | 188.40±2.30<sup>cd</sup> | 10.50±0.56<sup>ab</sup> |
| **T<sub>7</sub>** | 10.41±0.09<sup>bc</sup> | 5.60±0.06<sup>cd</sup> | 179.23±2.37<sup>e</sup> | 11.08±0.73<sup>a</sup> |
| **T<sub>8</sub>** | 10.11±0.28<sup>cd</sup> | 5.62±0.07<sup>cd</sup> | 200.17±0.84<sup>a</sup> | 11.17±0.57<sup>a</sup> |
| **T<sub>9</sub>** | 10.87±0.16<sup>a</sup> | 5.96±0.08<sup>b</sup> | 187.93±3.39<sup>cd</sup> | 11.42±0.62<sup>a</sup> |
| **T<sub>10</sub>** | 9.67±0.14<sup>e</sup> | 5.18±0.21<sup>ef</sup> | 177.33±1.82<sup>e</sup> | 9.83±0.90<sup>cde</sup> |

**T<sub>1</sub>:**100% RDF alone, **T<sub>2</sub>:** 75% RDF + soil micronutrients + one foliar spray of micronutrients, **T<sub>3</sub>:** 50% RDF + soil micronutrients + one foliar spray of micronutrients, **T<sub>4</sub>:** 25% RDF + soil micronutrients + one foliar spray of micronutrients, **T<sub>5</sub>:** 75% RDF + soil micronutrients + two foliar spray of micronutrients, **T<sub>6</sub>:** 50% RDF + soil micronutrients + two foliar spray of micronutrients, **T<sub>7</sub>:** 25% RDF + soil micronutrients + two foliar spray of micronutrients, **T<sub>8</sub>:** 75% RDF + two foliar spray of micronutrients, **T<sub>9</sub>:** 50% RDF + two foliar spray of micronutrients, **T<sub>10</sub>:** 25% RDF+ two foliar spray of micronutrients.

### 3.4. Available macro- and micronutrients in soil

Significant differences were observed in soil nutrient availability **(Table 5).** Available N was highest under 75% RDF + two foliar sprays, followed by integrated treatments receiving 50-75% RDF with micronutrients; the lowest N was found in 25% RDF + soil + foliar micronutrients. Available P was highest in 75% RDF + foliar micronutrients, with integrated treatments maintaining higher P than the control. Minimum P content occurred under 25% RDF + foliar micronutrients. Soil K content followed a similar pattern, with the highest values under 75% RDF + foliar spray and the lowest under 25% RDF + micronutrients. Micronutrient availability (Table 6) also varied significantly. Fe and Cu were highest in 50% RDF + soil + foliar micronutrients. Zn, B, and Ca were highest in 75% RDF + soil + foliar micronutrients, whereas Ca was lowest under 100% RDF.

**Table 5.**
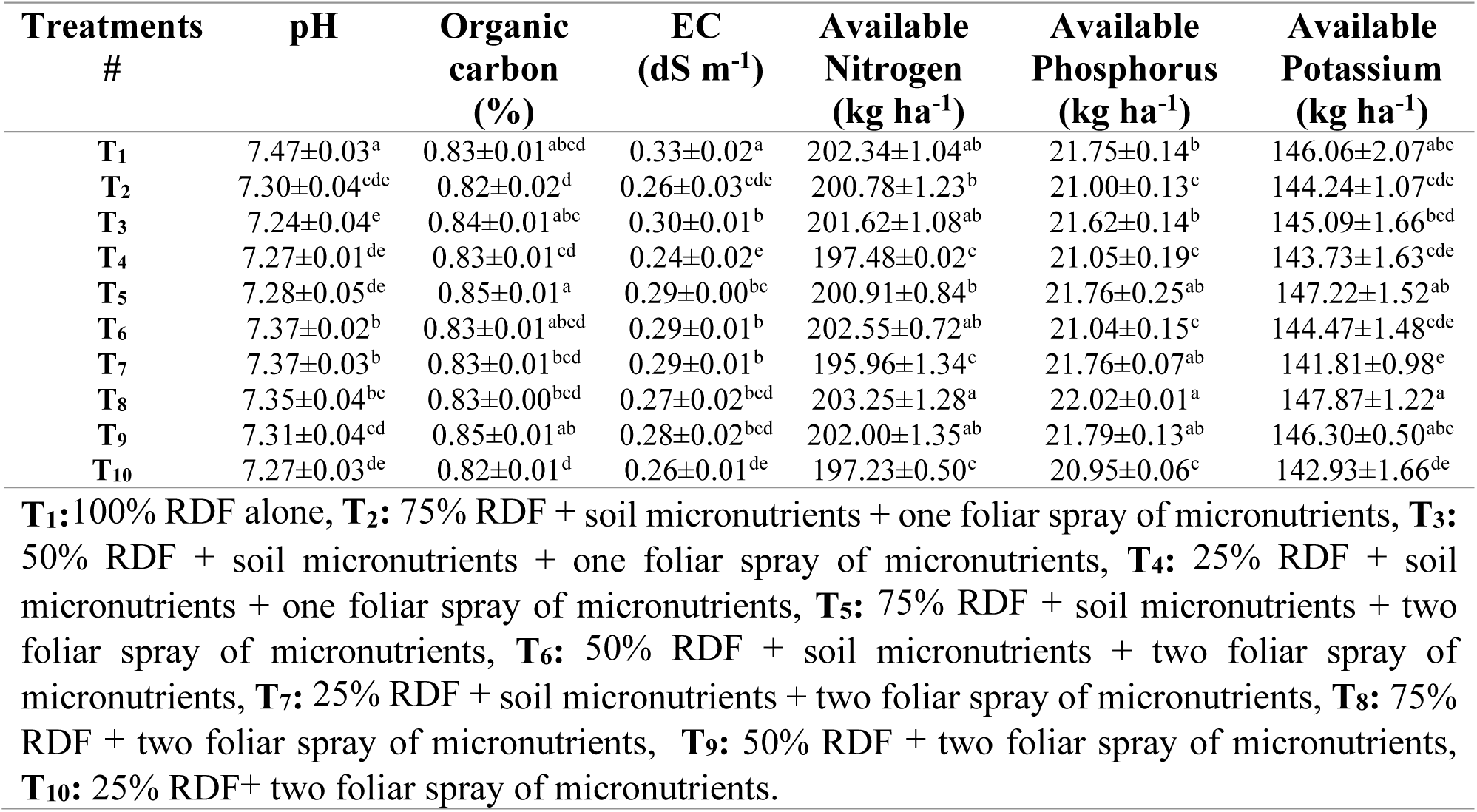
Influence of macro and micronutrients on soil properties (pH, organic carbon, EC (dS m^-1^), available nitrogen, phosphorus, and potassium) in a mango cv. Dashehari under medium density planting (**Mean of two years**).

**Table 6.**
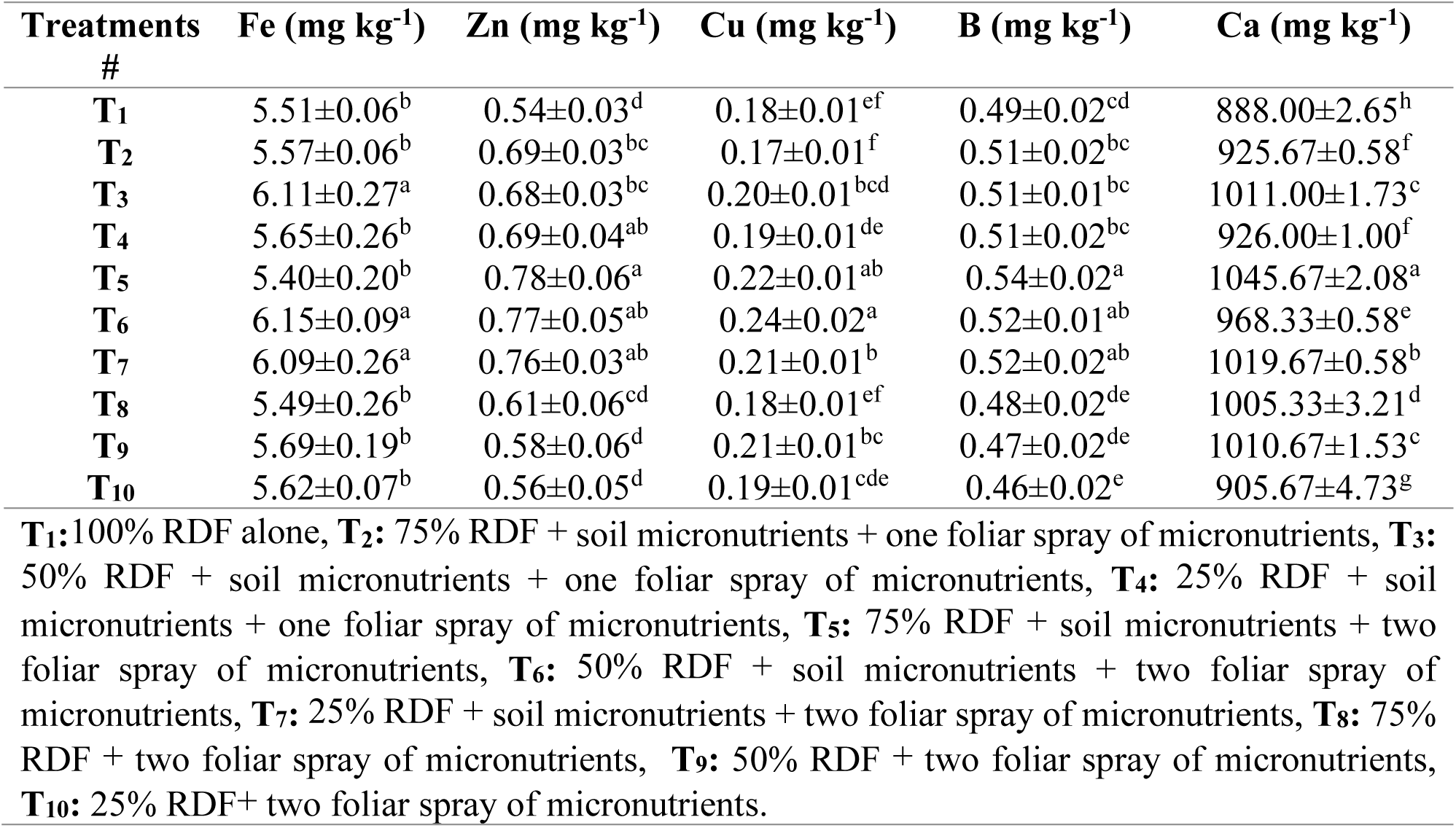
Influence of macro and micronutrients on Fe, Zn, Cu, B & Ca content of soil in mango cv. Dashehari under medium density planting (**Mean of two years**).

### 3.5. Leaf macro- and micronutrients

Leaf macronutrient concentrations varied significantly **(Fig. 7a-c).** Leaf N ranged from 0.93-1.07%, with the highest values under 75% RDF + two foliar sprays. Leaf P ranged 0.074-0.130%, with the maximum under 75% RDF + foliar spray. Leaf K ranged 0.50-0.60%, highest in 75% RDF + foliar spray. The lowest N, P and K were recorded in reduced RDF with limited micronutrient inputs. Leaf micronutrients (Fig. 8a–d) also differed significantly. Fe ranged 77.53-80.70 ppm, Zn 15.78-19.45 ppm, Cu 8.97-12.20 ppm, B 45.76-54.03 ppm, and Ca 1.38-1.86%. The highest values for most micronutrients occurred under 75% RDF + foliar sprays, while reduced RDF with foliar application alone produced the lowest concentrations.

**Figure 7.**
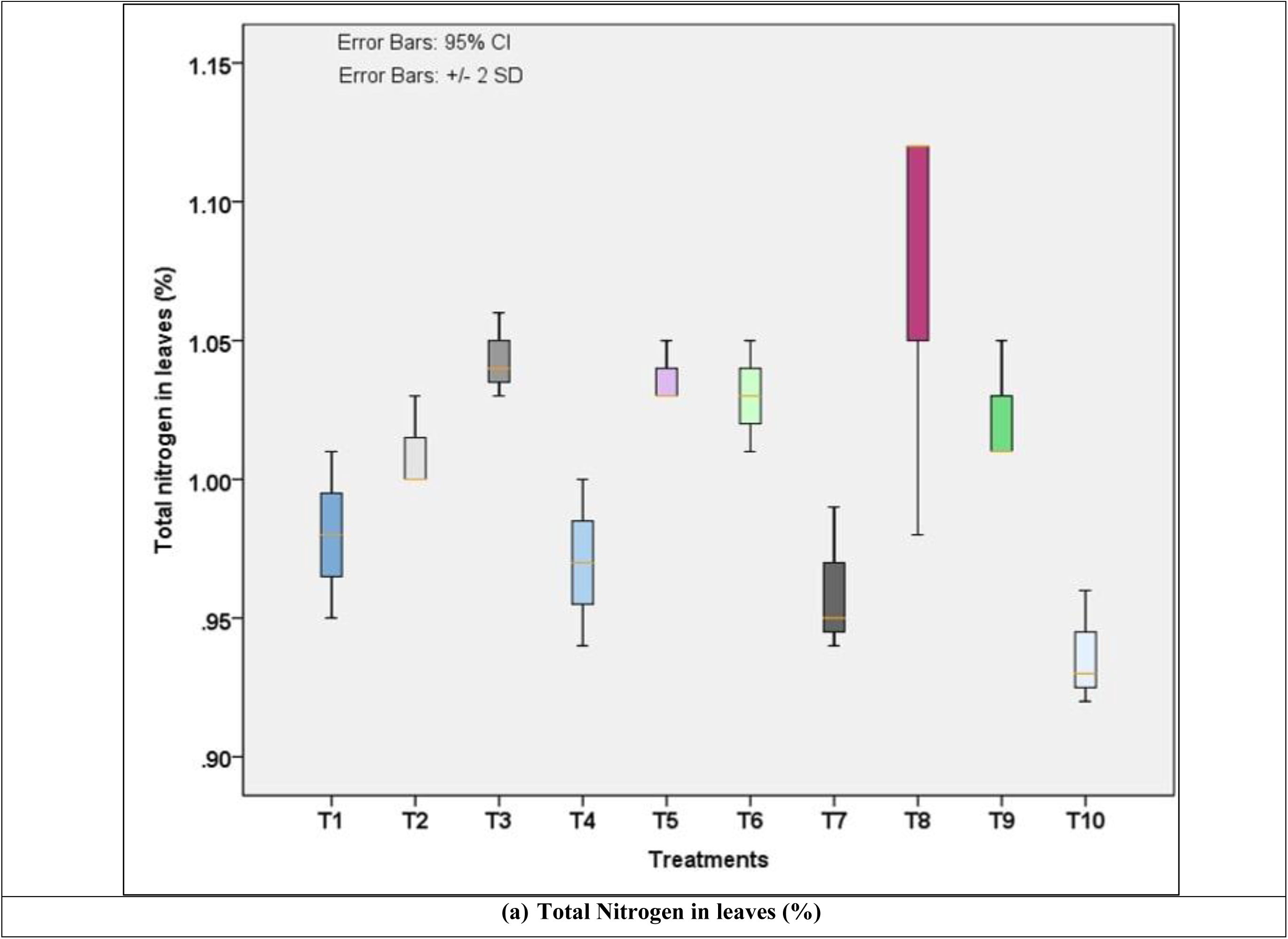
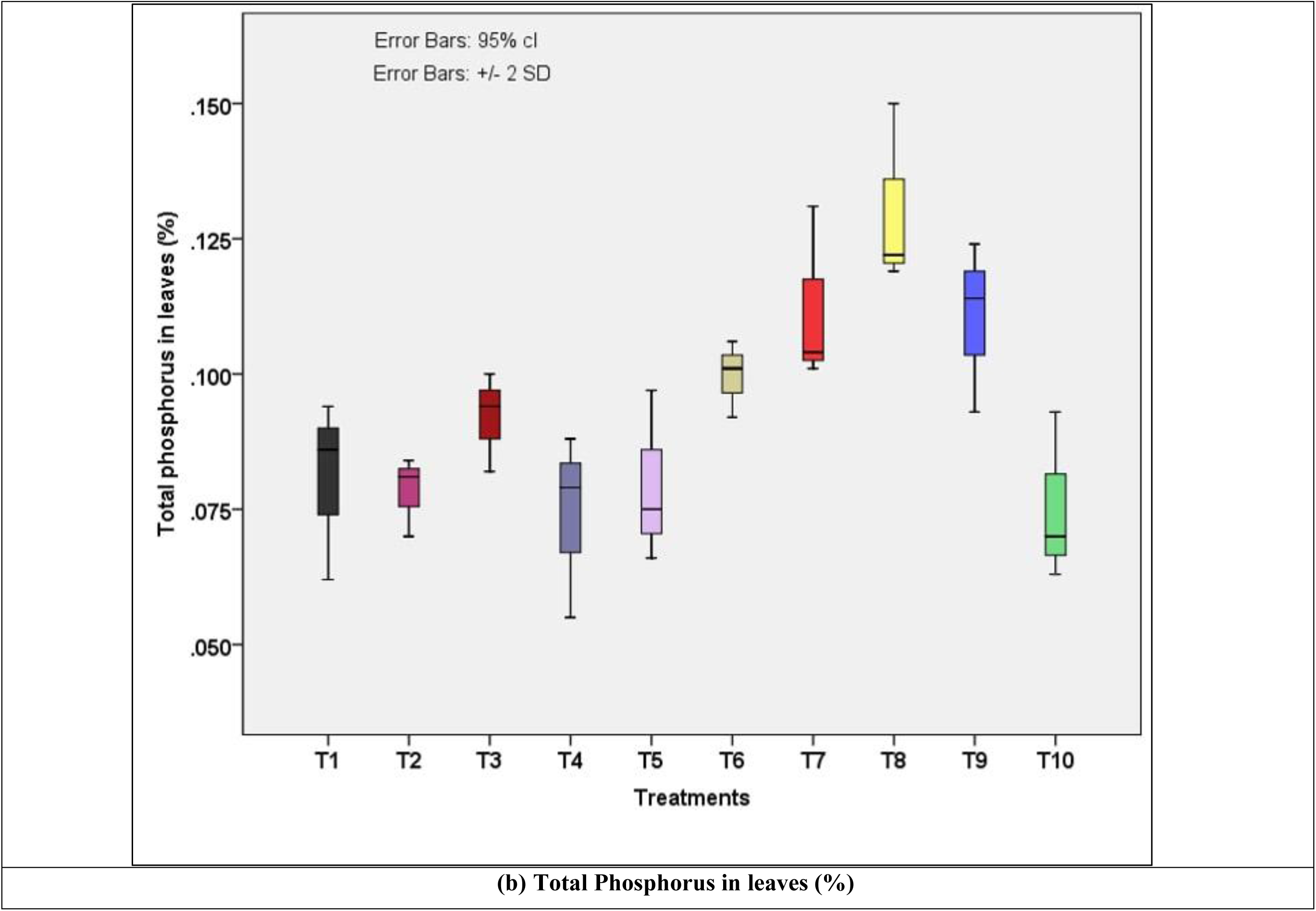
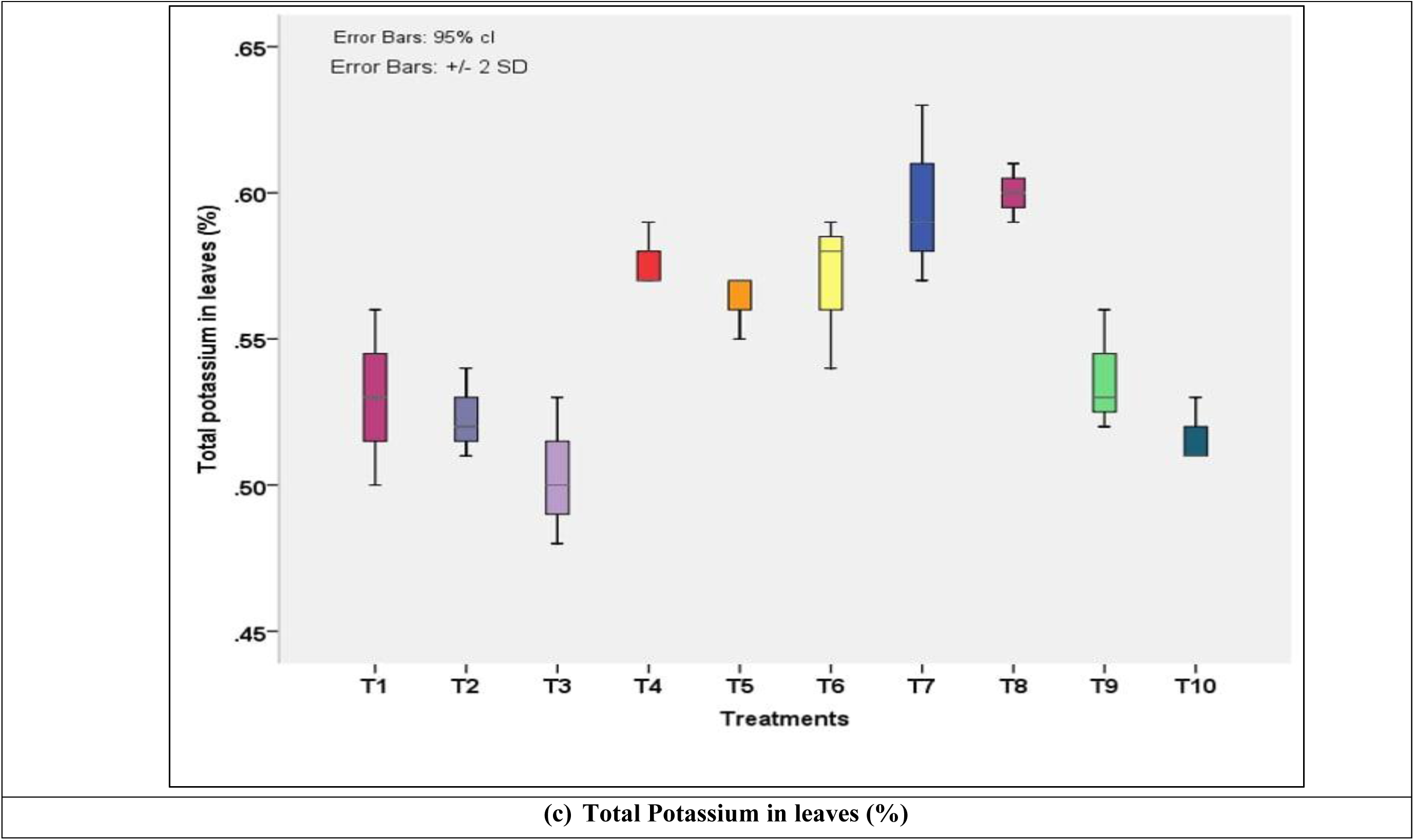
Influence of INM practices on leaf macronutrient (a) nitrogen, (b) phosphorus, and (c) Potassium content of mango (*Mangifera indica* L.) cv. Dashehari under medium-density planting (Mean of two years). T1:100% RDF alone, T2: 75% RDF + soil micronutrients + one foliar spray of micronutrients, T3: 50% RDF + soil micronutrients + one foliar spray of micronutrients, T4: 25% RDF + soil micronutrients + one foliar spray of micronutrients, T5: 75% RDF + soil micronutrients + two foliar spray of micronutrients, T6: 50% RDF + soil micronutrients + two foliar spray of micronutrients, T7: 25% RDF + soil micronutrients + two foliar spray of micronutrients, T8: 75% RDF + two foliar spray of micronutrients, T9: 50% RDF + two foliar spray of micronutrients, T10: 25% RDF+ two foliar spray of micronutrients.

**Figure 8.**
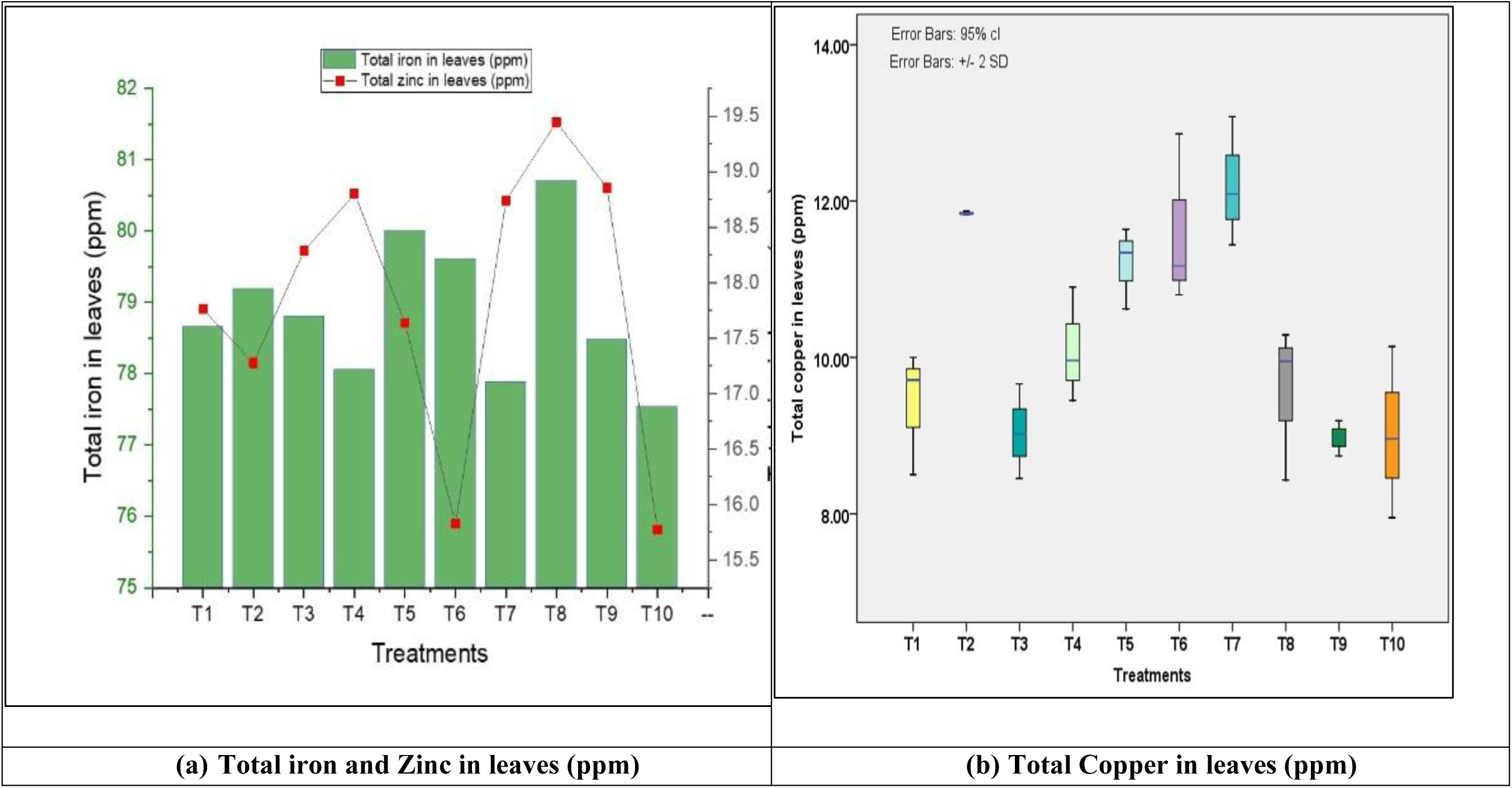
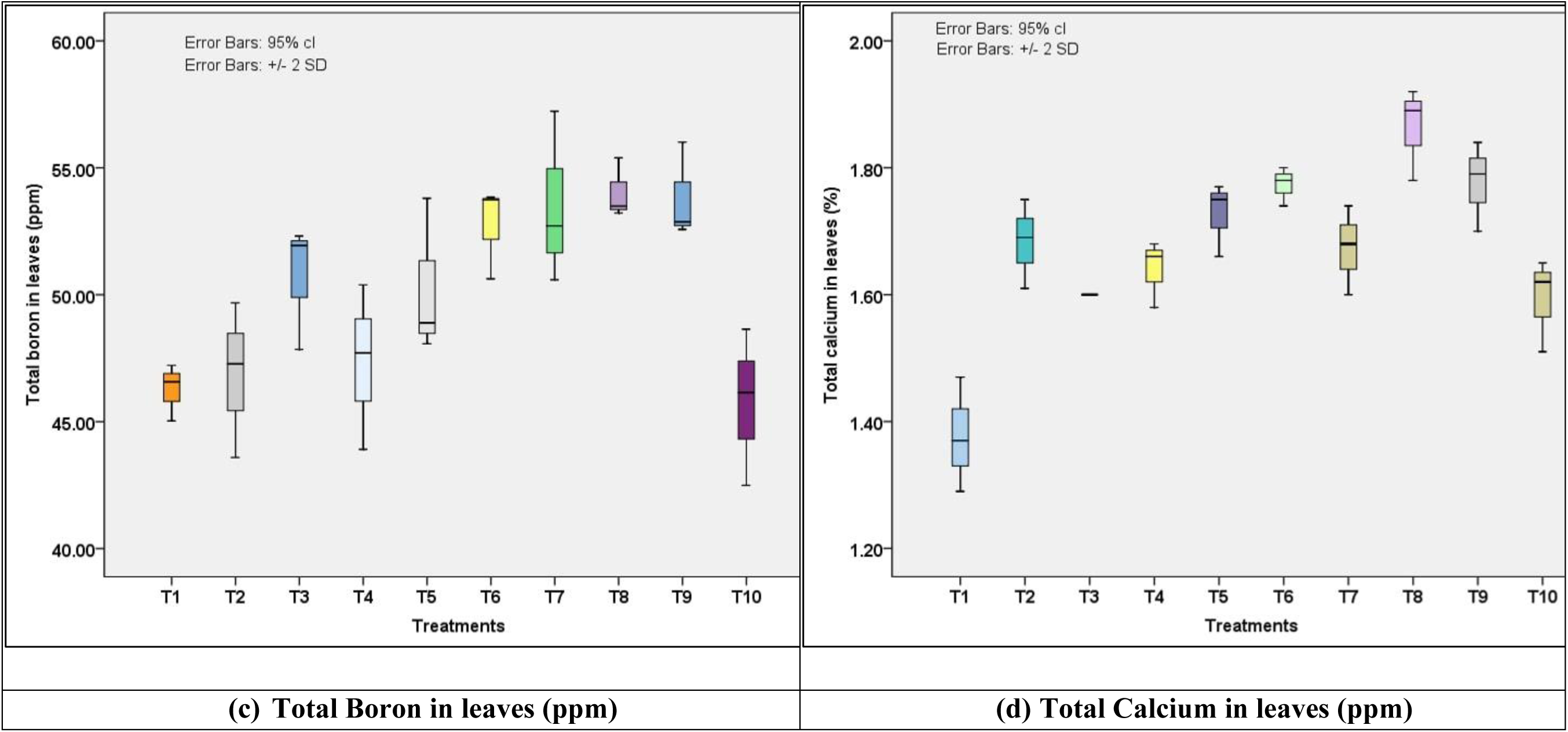
Influence of INM practices on leaf micronutrient (a) iron & zinc (b) copper (c) boron (d) calcium content of mango (*Mangifera indica* L.) cv. Dashehari under medium-density planting (Mean of two years). T1:100% RDF alone, T2: 75% RDF + soil micronutrients + one foliar spray of micronutrients, T3: 50% RDF + soil micronutrients + one foliar spray of micronutrients, T4: 25% RDF + soil micronutrients + one foliar spray of micronutrients, T5: 75% RDF + soil micronutrients + two foliar spray of micronutrients, T6: 50% RDF + soil micronutrients + two foliar spray of micronutrients, T7: 25% RDF + soil micronutrients + two foliar spray of micronutrients, T8: 75% RDF + two foliar spray of micronutrients, T9: 50% RDF + two foliar spray of micronutrients, T10: 25% RDF+ two foliar spray of micronutrients.

### 3.6. Path coefficient analysis

PLS-SEM path coefficient analysis **(Fig. 9)** showed that plant nutrients exerted the highest direct positive effect on fruit yield (0.323), followed by plant growth (0.311) and soil nutrients (0.310).

**Figure 9.**
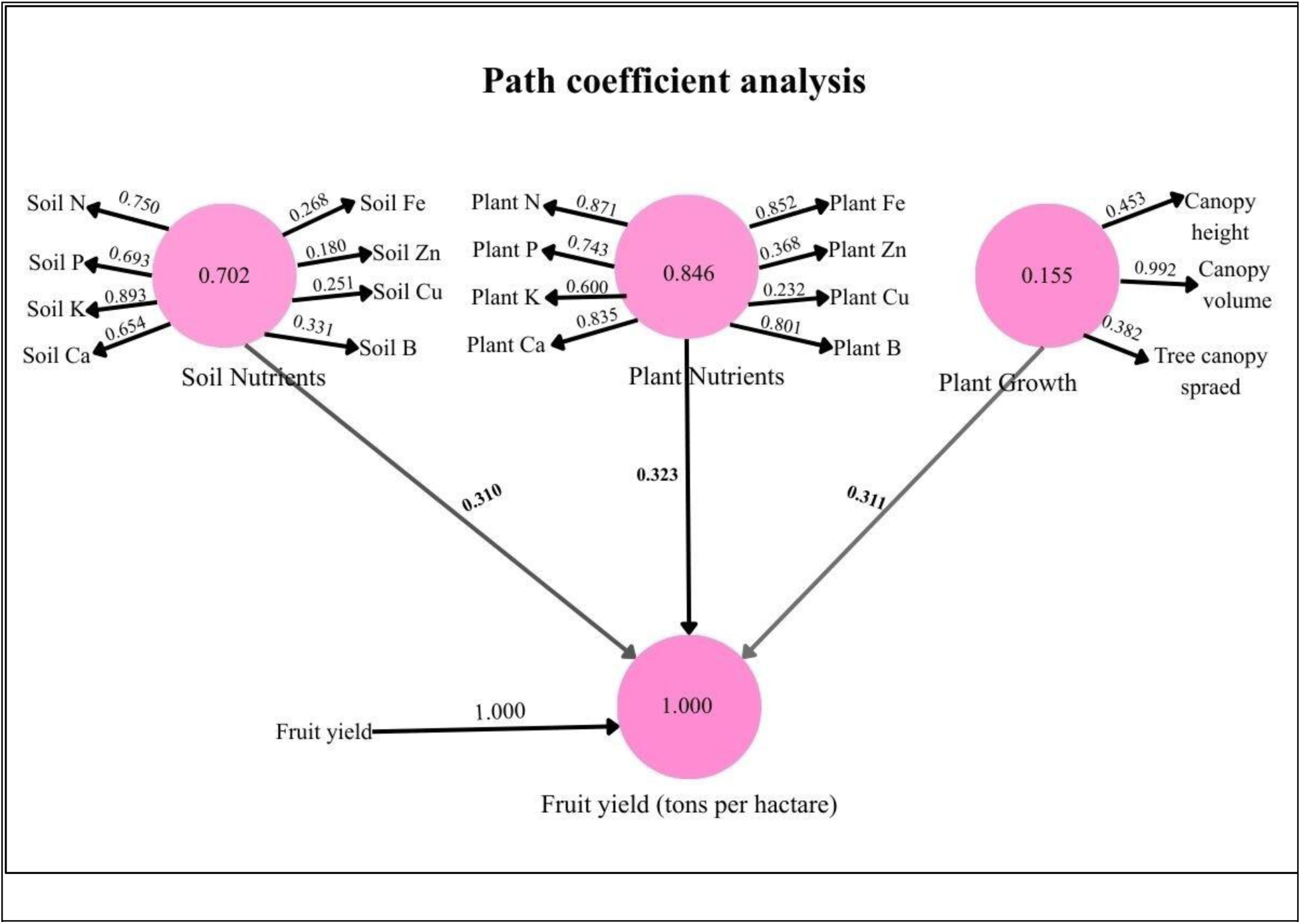
Path diagram showing direct and indirect effects of soil nutrients, plant nutrients, and plant growth on fruit yield of mango (*Mangifera indica* L.) cv. Dashehari under medium-density planting (Mean of two years). T1:100% RDF alone, **T2:** 75% RDF + soil micronutrients + one foliar spray of micronutrients, **T3:** 50% RDF + soil micronutrients + one foliar spray of micronutrients, **T4:** 25% RDF + soil micronutrients + one foliar spray of micronutrients, **T5:** 75% RDF + soil micronutrients + two foliar spray of micronutrients, **T6:** 50% RDF + soil micronutrients + two foliar spray of micronutrients, **T7:** 25% RDF + soil micronutrients + two foliar spray of micronutrients, **T8:** 75% RDF + two foliar spray of micronutrients, **T9:** 50% RDF + two foliar spray of micronutrients, **T10:** 25% RDF+ two foliar spray of micronutrients.

Within the soil nutrient construct, macronutrients contributed strongly, with the highest loadings for soil K (0.893), soil N (0.750) and soil P (0.693). Micronutrients showed lower loadings (Fe 0.268, Zn 0.180, Cu 0.251). Within the plant nutrient construct, leaf N (0.871), Ca (0.835) and Fe (0.852) showed the highest loadings. Canopy volume had the highest loading (0.992) within the plant growth construct.

### 3.7. Principal component analysis

PCA identified distinct grouping patterns associated with nutrient management treatments **(Fig. 10).** PC1 and PC2 explained 91.85% of total variability, with PC1 contributing 79.25%. Fruit yield (kg plant⁻¹ and t ha⁻¹) and percent increase in canopy volume loaded strongly on PC1. Fruit retention, percent fruit drop and canopy spread were associated with PC2. Soil N, soil K and leaf P clustered along PC1. The treatment 75% RDF + two foliar sprays aligned most closely with yield- and canopy-related vectors. Treatments receiving 75% or 50% RDF + soil + foliar micronutrients (T5, T7, T9) clustered near major growth and yield attributes. Reduced RDF with limited micronutrients (T1, T2, T3, T4, T6, T10) formed separate clusters.

**Figure 10.**
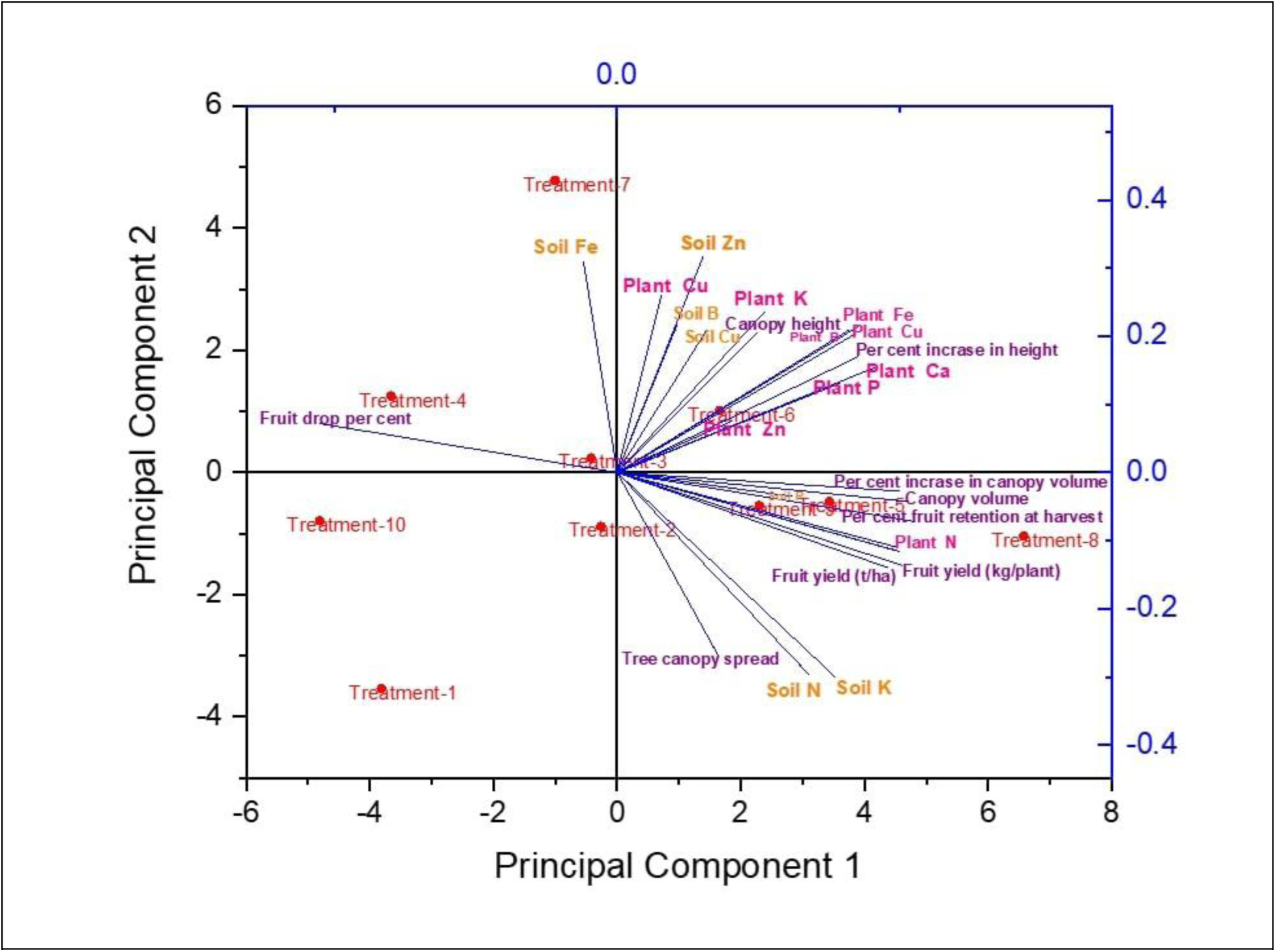
PCA biplot showing the relationship of INM treatments with soil nutrients, leaf nutrient status, growth, yield, and fruiting attributes of mango (*Mangifera indica* L.) cv. Dashehari under medium-density planting. T1:100% RDF alone, **T2:** 75% RDF + soil micronutrients + one foliar spray of micronutrients, **T3:** 50% RDF + soil micronutrients + one foliar spray of micronutrients, **T4:** 25% RDF + soil micronutrients + one foliar spray of micronutrients, **T5:** 75% RDF + soil micronutrients + two foliar spray of micronutrients, **T6:** 50% RDF + soil micronutrients + two foliar spray of micronutrients, **T7:** 25% RDF + soil micronutrients + two foliar spray of micronutrients, **T8:** 75% RDF + two foliar spray of micronutrients, **T9:** 50% RDF + two foliar spray of micronutrients, **T10:** 25% RDF+ two foliar spray of micronutrients.

### 3.8. Pearson correlation coefficients

Pearson correlation analysis **(Table 7)** showed significant associations among yield, growth, and nutrient variables. Fruit yield correlated positively with fruit weight (r=0.58, P<0.01), soil P (r=0.58, P<0.01), soil Zn (r=0.48, P<0.05), and shelf life (r=0.49, P<0.05). Canopy volume correlated with fruit yield (r=0.41) and fruit weight (r=0.52, P<0.01). Fruit weight correlated with canopy volume (r=0.52, P<0.01), leaf Fe (r=0.48, P<0.05), soil Zn (r=0.52, P<0.01), and soil P (r=0.45, P<0.05).

**Table 7.**
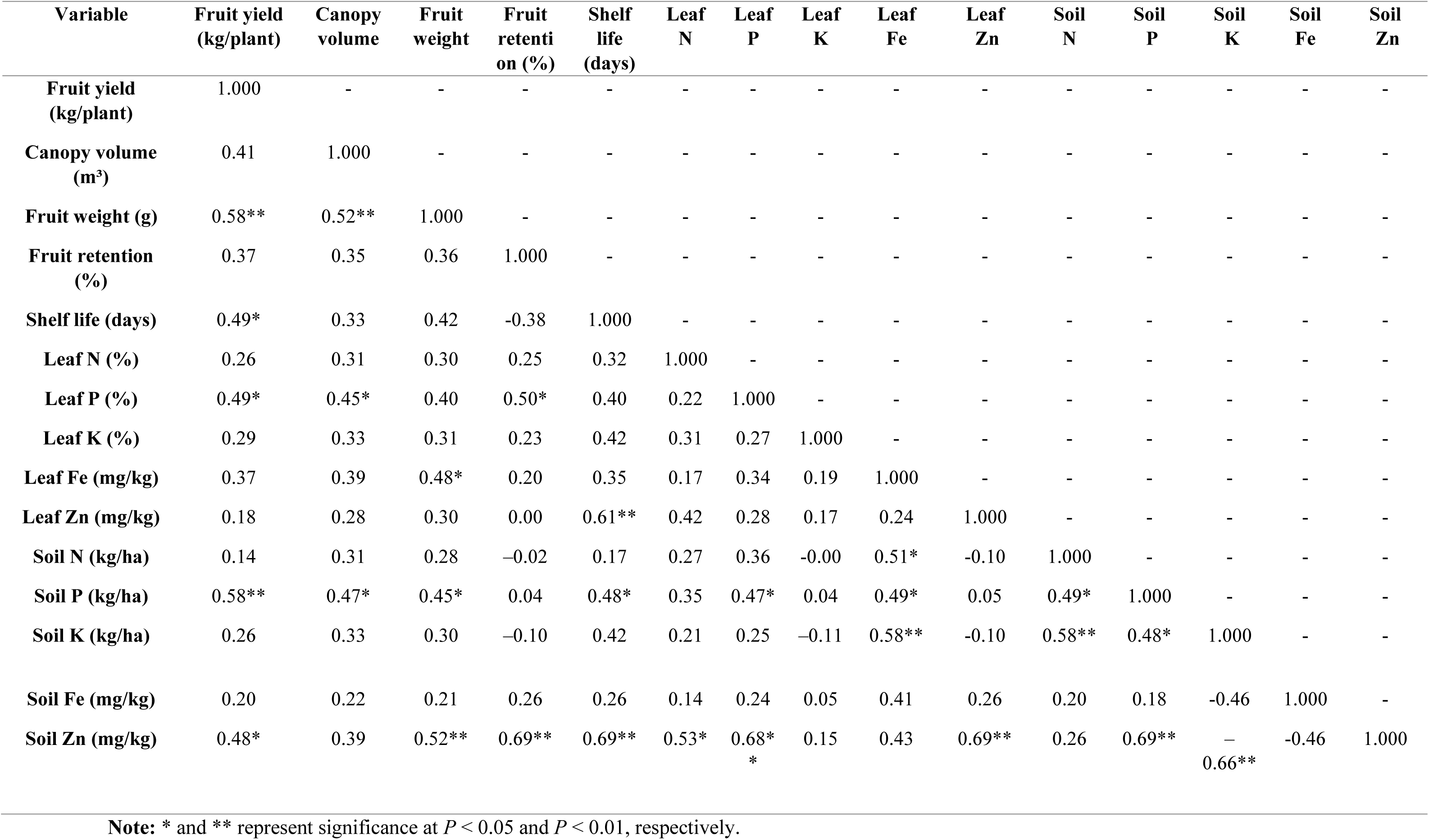
Pearson correlation coefficients among fruit yield, growth, soil, and plant nutrient variables in medium-density mango orchards.

Fruit retention showed strong correlations with soil Zn (r=0.69, P<0.01) and leaf P (r=0.50, P<0.05). Shelf life correlated with leaf Zn (r=0.61, P<0.01), soil Zn (r=0.69, P<0.01), and soil P (r=0.48, P<0.05). Leaf P correlated with canopy volume (r=0.45, P<0.05), fruit retention (r=0.50, P<0.05), and soil P (r=0.47, P<0.05). Soil Zn showed strong associations with fruit retention, shelf life, fruit weight, leaf N, leaf P, and soil P, and a negative correlation with soil K (r=–0.66, P<0.01).

## 4. Discussion

### 4.1. Vegetative growth

Integrated soil + foliar micronutrient treatments (notably 75% RDF + two foliar sprays and 50% RDF + soil + foliar) produced larger canopy height, spread, and volume than sole 100% RDF or reduced-RDF treatments with limited micronutrients. Foliar-applied Fe, Zn, B and Ca supplied readily available elements to active tissues, stimulating chlorophyll synthesis, auxin metabolism, cell division/elongation and lateral shoot formation; soil applications supported root uptake. These complementary pathways likely corrected latent deficiencies induced by prolonged NPK use and improved photosynthetic capacity and assimilate partitioning, promoting canopy expansion (Davenport et al. 2000; Normanly 1997; Xu et al. 2023, Lingwan et al., 2024). Boron’s role in cell-wall integrity and assimilate transport further supports vegetative expansion (Kohli et al. 2023; Ganie et al. 2013). Improved lateral shoot formation under integrated treatments indicates rapid correction of deficiencies and enhanced photosynthetic efficiency (Ullah et al. 2024; Khalil et al. 2025, Yadav et al., 2020). Canopy architecture, particularly volume, integrates plant nutrient status and light interception. Enhanced canopy under integrated treatments implies greater source strength for reproductive development while using reduced macronutrient inputs, supporting more sustainable fertilization strategies. Findings align with reports that micronutrient enrichment improves vegetative vigor in fruit crops and cereals, and with studies showing that exclusive NPK use can induce latent micronutrient limitations that restrict biomass accumulation (Srivastava & Hu 2019; Mohanasundaram et al., 2021; Balban et al. 2025; Khalil et al. 2025; Ullah et al. 2024).

### 4.2. Yield attributes, shelf life and B:C ratio

Integrated treatments increased fruits per panicle, fruit retention, fruit size and yield (highest under 75% RDF + two foliar sprays). Shelf life and B:C ratio improved in foliar-spray treatments; reduced-RDF alone (25% RDF) produced the lowest yields. Zn, B, Fe and Ca are vital for pollen viability, stigma receptivity, membrane integrity, and cell-wall stability, reducing physiological fruit drop and supporting successful fertilization (Álvarez-Herrera et al. 2025; Muengkaew et al. 2018). Calcium and boron also delay senescence and maintain firmness via effects on ethylene metabolism and membrane stability, extending shelf life. Improved internal nutrient status enhanced assimilate partitioning to fruit, increasing fruit weight. Supplementing reduced RDF with targeted micronutrients can sustain or enhance yield and postharvest quality while improving nutrient-use efficiency and profitability, offering a practical tool for sustainable mango production (Ravikiran 2018; Chaudhari & Singh 2019, Saini et al., 2020). Results corroborate studies reporting micronutrient benefits for fruit set, retention, and postharvest quality (Sankar et al. 2013; Muengkaew et al. 2018; Bhardwaj et al. 2022) and extend them by demonstrating that integrated soil + foliar approaches under reduced RDF maintain productivity and economics better than single-source nutrient strategies.

### 4.3. Soil pH, organic carbon, and electrical conductivity

Soil pH was higher under continuous 100% RDF; integrated micronutrient treatments produced slightly lower pH. Organic carbon (OC) varied minimally but tended to be higher in integrated treatments; EC was highest under continuous 100% RDF and moderated under integrated management. Repeated inorganic fertilization alters ionic balance and may increase exchangeable basic cations, raising pH (Pahalvi et al. 2021; Ganeshamurthy et al. 2016, Lingwan et al.,2025). Sulphate forms of micronutrients and enhanced root exudation can acidify the rhizosphere (Sharma et al. 2025; Houmani et al. 2015; Rengel 2015). Balanced nutrition and improved microbial activity under integrated management can slightly increase OC and stabilize carbon pools (Bhatt et al. 2025; Dhaliwal et al. 2024). Lower EC under integrated regimes reflects reduced soluble salt accumulation when foliar supply substitutes part of soil-applied inputs (Chhabra 2022; Job et al., 2022; Kaya & Ashraf 2024; Srivastava & Hota 2025). Integrating micronutrients with reduced RDF may help stabilize soil reaction, sustain OC and limit salinity build-up advantages for long-term soil health. Observations are consistent with literature documenting fertilizer-driven changes in soil pH and EC and benefits of integrated nutrient regimes in moderating adverse chemical trends (Kailasrao 2016; Marschner 2012, Job et al., 2025; Kushwaha et al., 2022).

### 4.4. Available soil macro- and micronutrients

Integrated treatments (especially 75% RDF + foliar) maintained higher available N, P and K than reduced-RDF alone. Micronutrient availability patterns varied by treatment (e.g., Fe and Cu high in 50% RDF + soil + foliar; Zn, B, Ca high in 75% RDF + soil + foliar). Foliar feeding reduces immediate soil withdrawal; soil micronutrients and improved root function support mineralization and microbial processes, maintaining available pools. Micronutrients can modulate P solubilization and microbial enzyme activity, conserving soil P and K (Dhaliwal et al. 2024; Rajput 2019; Hu et al. 2023, Ali et al., 2025). Integrated nutrient management synchronizes soil nutrient pools with crop demand, improving nutrient-use efficiency and reducing depletion risks associated with unbalanced fertilization. Results align with prior studies showing that combined soil–foliar strategies sustain macro- and micronutrient availability better than reliance on macronutrients alone (Niu et al. 2021; Maity et al. 2021).

### 4.5. Plant nutrient status

Leaf N, P, K, and micronutrient concentrations were highest under integrated treatments (leaf N 0.93-1.07%; Fe 77.5-80.7 ppm), with the lowest values in reduced-RDF with minimal micronutrients. Foliar supply increases tissue concentrations quickly; enhanced root functioning improves soil uptake and translocation. Micronutrients facilitate enzymes and transporters that improve macronutrient assimilation (Pilli et al. 2025; Ahmad et al. 2018; Singh et al. 2022, Ali & Lingwan 2025). Leaf nutrient status better reflects internal availability and predicts yield more closely than soil pools (see path analysis). Routine leaf diagnostics can guide targeted foliar/soil applications. Findings corroborate studies showing immediate tissue enrichment after foliar feeding and superior plant nutrition under integrated approaches (Sun et al., 2021; Khan & Ahmed 2021; Thakur et al.,2025).

### 4.6. Systems analyses (PLS-SEM, PCA, correlations)

PLS-SEM indicated plant nutrients had the strongest direct effect on yield (path coefficient = 0.323), followed by plant growth (0.311) and soil nutrients (0.310). PCA clustered high-performing integrated treatments with yield and canopy vectors; Pearson correlations linked yield closely to fruit weight, soil P, and soil Zn. Integrated management improved internal nutrient pools and canopy architecture, which together regulate source-sink dynamics central to yield. Macronutrients form the metabolic base while micronutrients modulate physiological efficiency, explaining the observed factor loadings. Management should target tissue nutrition and canopy maintenance in addition to soil fertility. Foliar diagnostics and targeted sprays that improve plant nutrient status will be effective levers to increase yield. Structural and multivariate outcomes are consistent with prior research emphasizing internal plant nutrition and canopy traits as proximal drivers of productivity (Sun 2020; Kishore et al. 2023; Bukhari et al. 2025).

## Conclusion

This study demonstrates that integrated nutrient management, particularly the combination of 75% RDF with two foliar micronutrient sprays, is more effective than the conventional full RDF approach for sustaining growth, nutrient balance, and productivity in medium-density mango orchards. Integrated soil-foliar micronutrient supplementation improved canopy development, enhanced fruit set and retention, increased fruit size and yield, and extended shelf life. These improvements were accompanied by higher soil and leaf concentrations of macro- and micronutrients, indicating more efficient nutrient uptake and physiological utilization. Multivariate analyses (PCA and PLS-SEM) further revealed that plant nutrient status is the strongest determinant of yield, followed by canopy attributes and soil nutrient pools. The alignment of high-performing treatments with yield-related variables underscores the role of balanced nutrient dynamics in supporting source-sink functioning in perennial fruit systems. Importantly, reducing the RDF to 75% while supplying targeted foliar micronutrients improved nutrient-use efficiency and economic returns, offering a cost-effective and environmentally sustainable alternative to full-dose chemical fertilization. In the future, more effective high-throughput approaches and systems biology can be adopted to enhance in-depth understanding of the physiology and biochemistry under different nutrient media ( Koley et al., 2024, Jaroensuk et al.,2025). Overall, the findings highlight that 75% RDF + foliar micronutrient sprays are a viable, resource-efficient strategy for enhancing productivity, soil health, and profitability in mango orchards under medium-density planting. This approach provides a scientifically validated, site-specific framework that can support long-term sustainability and resilience in mango cultivation.

## Author’s contribution

Kuldeep: Data collection, Formal analysis, Investigation, Methodology, Validation, Writing - draft, and execution of the entire research work.

Ashok Kumar Singh: Survey, Data curation, Writing-draft, review & editing.

Amit Bhatnagar, Omveer Singh, Shailesh Chandra Shankhdhar, Satya Pratap Pachauri, Manoj Kumar Bhatt: Conceptualization of the work thought, Software, Data analysis/interpretation, Writing - review & editing.

## Acknowledgment

Acknowledgment is extended for the financial assistance and infrastructural support provided by the Horticulture Research Centre, Patharchatta, and the Department of Horticulture, GBPUA&T, Pantnagar, Uttarakhand, India.

